# Method to quickly map multifocal Pupillary Response Fields (mPRF) using Frequency Tagging

**DOI:** 10.1101/2023.06.28.546061

**Authors:** Jean Lorenceau

## Abstract

We present a method for mapping multifocal Pupillary Response Fields in a short amount of time, using a visual stimulus covering 40° of visual angle, divided in 9 contiguous sectors, simultaneously modulated in luminance at specific, incommensurate, temporal frequencies. We tested this multiple Pupillary Frequency Tagging (mPFT) approach with young healthy participants (N=36), and show that the spectral power of the sustained pupillary response elicited by 45 seconds of fixation of this multipartite stimulus reflects the relative contribution of each sector/frequency to the overall pupillary response. We further analyze the phase lag for each temporal frequency as well as several global features related to pupil state. Test retest performed on a subset of participants indicates good repeatability. We also investigate the existence of structural (RNFL)/functional (mPFT) relationships. We then summarize results of clinical studies conducted with mPFT on patients with neuropathies and retinopathies and show that the features derived from pupillary signal analyzes, the distribution of spectral power in particular, allows sorting patients from healthy participants with excellent sensitivity and specificity. This method thus appears a convenient, objective and fast tool for assessing the integrity of retino-pupillary circuits, as well as idiosyncrasies, that permits to objectively detect or follow-up retinopathies or neuropathies in a short amount of time.

## Introduction

Renewed interest in pupillary responses manifests by a tremendous increase of publications in recent years. Owing to the particular circuits underlying pupillary dilation and constriction (Szabadi and Bradshaw, 1996; Wang and Munoz, 2015), and to the discovery of intrinsically responsive retinal ganglion cells (ipRGCs containing melanopsin, Provencio et al., 1999; Hattar et al, 2002), two distinct fields, rooted in the seminal studies conducted in the sixties (e.g. Kahneman & Beatty, 1966), developed, thanks to the dissemination of eye-tracking technologies. Publications are either concerned by pupillary dilation – mydriasis, mainly related to a sympathetic drive associated to cognitive functions-, or to pupillary constriction elicited by light –myosis, mainly involving the parasympathetic pathway- and mostly used to probe Eye Health. It is out of the scope of the present study to summarize the very numerous publications related to both fields, as several recent reviews are available to appreciate this renewed interest (e.g. WilHelm & Wilhelm, 2003; Wang and Munoz, 2015; Mathot, 2018; Zele and Gamlin, 2020).

In both the “Clinical” and “Cognitive” fields, with some exceptions (Najjar et al., 2021), the focus is on transient pupillary responses, either elicited by flashes of light (Pupil Light Reflex, PLR), or by timely controlled –cognitive-tasks (involving attention, memory, decision, confidence, etc.).

In this dynamic context, we searched for novel, convenient and fast ways of assessing pupillary reactivity to light in different regions of the visual field.

Our aims are twofold:

1. to provide a tool able to characterize the idiosyncratic heterogeneity of pupillary responses in the visual field of healthy individuals, which could help to understand and to account for inter-individual differences in visuo-cognitive tasks,
2. to use pupillary responses to spatially distributed light stimulation to objectively assess visual field defects in ophthalmic –or neurologic-diseases, so as to complement the functional examinations (Standard Automated Perimetry in particular), that rely on subjective responses, are long (> 7 min) and tedious. The rationale is that a retinal defect in a particular location, related to neuropathies or retinopathies, should alter the pupillary responses in that same location.

We here present a method relying on sustained multiple pupillary frequency tagging (mPFT). We found that up to 9 temporal frequencies associated to 9 non-overlapping regions could be mixed while still eliciting pupillary responses whose spectral power reflects the contribution of each frequency/region. With this approach, a “Pupillary Response Field” can be obtained at once in less than a minute. The fact that a “Pupillary Response Field” is assessed simultaneously in several regions of the visual field is an important and interesting feature of mPFT: it permits to compare the *relative* contribution of each region which should be immune to sequential effects that could bias the responses obtained at different times, as can be the case with sequentially presented light flashes, or to the comorbidities or medication that may alter the global pupillary responses. In addition, using a continuous stimulation avoids the need to return to baseline in between flashes, as is often the case, such that all the recorded data are relevant. Finally, using continuous recordings permit to assess additional relevant variables, as for instance the eye-movements or the number of blinks made during sustained fixation, known to be altered in some ophthalmic pathologies.

We present below the stimulus and methodology that we tested with 36 young healthy adults, detail the analysis pipeline we developed, describe the results obtained with this method, and discuss some limitations and advantages of mPFT. We further analyze the correlation of pupillary spectral power with structural data (RNFL). We end with a summary of clinical studies (Ajasse et al., 2022; Stelandre et al., 2023; Trinquet et al, in preparation) performed with mPFT on several diseases (neuropathies and retinopathies) that demonstrate that this method provide excellent classification results, opening the way to be routinely used in clinical settings.

## Method and Stimuli

### Stimulus

The stimulus consists in 9 sectors of different sizes, overall covering about 40° of visual angle, arranged so as to stimulate central, paracentral and peripheral regions (Figure 1). The sizes of the peripheral, paracentral and central sectors were chosen on the basis of extensive preliminary experiments (Ajasse, 2019) so as to approximatively match the retinal magnification factor. These sizes may however be adapted (see discussion below).

**Figure 1:**
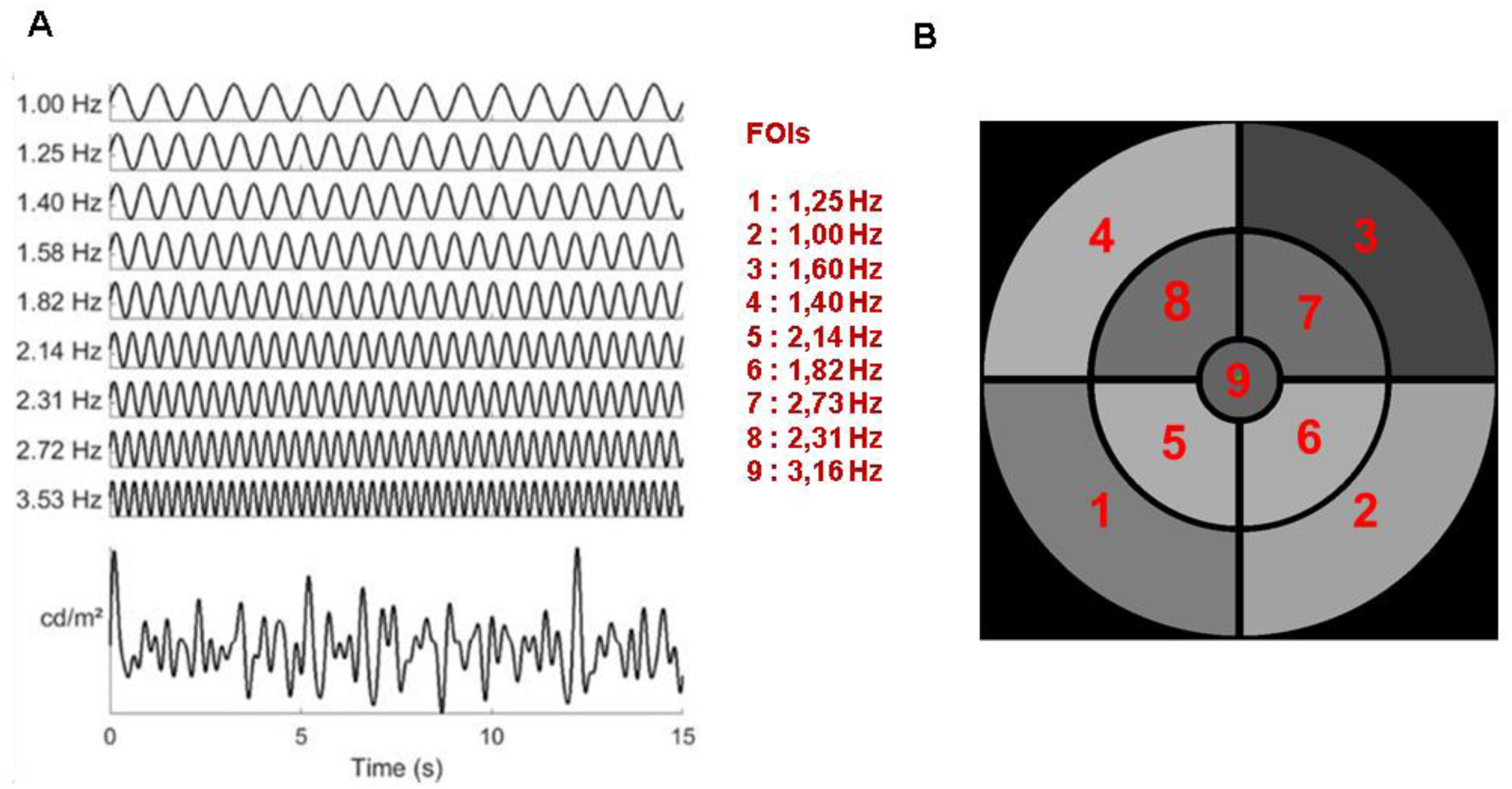
**A:** Distribution of the temporal modulation frequencies (TMFs) and the resulting overall luminance modulation. **B.** Stimulus configuration of the 9 sectors, each coupled with a TMF denoted by its index. The stimulus subtends about 40° of visual angle at 57 cm (central disk 4.6°; paracentral sectors, 5-19.6°; Peripheral sectors 20-40°). See Movie #1.

Each sector is periodically modulated in luminance at a specific temporal modulation frequency (TMF), different for each sector and incommensurate with the other TMFs. Supplementary Movie #1 shows an excerpt of the stimulus used in the study.

The temporal transfer function of the pupil, as assessed by Clarke et al (2003) in non-human primates, is low pass, with an upper limit of about 5 Hz, constraining the choice of the temporal frequencies that can be used to tag pupillary activity. The choice of the TMFs was dictated by several considerations: 1. the highest TMF in the stimulus must be lower than 4 Hz to ensure that reliable and sustained pupil responses would be elicited during stimulation. 2. The lowest TMF should be around 1 Hz, to ensure that a too limited number of cycles during a run would not bias spectral analyzes. 3. TMFs were chosen to be incommensurate, so as to avoid that harmonics and fundamental frequencies overlap, as this could introduce artifacts and alter the analyses of the pupil responses. In addition, care was taken to avoid as much as possible FOIs corresponding to intermodulation products, equal to the sum and difference between frequencies that may appear in the Fourier spectrum if non-linear interactions between frequencies would occur. The 4 lowest TMFs were associated to the 4 eccentric annulus sectors; the 4 intermediate TMFs were associated to the para-central annulus sectors. The highest TMF was presented in central vision to weight the, otherwise dominant, foveal contribution to pupillary activity (Hong e al, 2001). As a result of these constraints, the chosen TMFs were: 1.00 1.25 1.39 1.58 1.81 2.14 2.31 2.73 and 3.16 Hz (Figure 1). The minimum duration and the minimum amplitude of the sustained stimulation that still evoked reliable pupillary responses at all Frequencies of Interest (FOIs) were assessed in preliminary experiments (Ajasse, 2019). The shortest stimulation duration we found was about 45 sec. The mean luminance of each sector, expressed in RGB units, was set to 127 (Red, Green and Blue guns set to 127) and the modulation amplitude to 60 (in RGB units). With the screen settings used for that study, this corresponds to a mean luminance of 51 cd/m², a maximum luminance of 100 cd/m² and a minimum luminance of 20 cd/m². Note that the same stimulus was employed for the left and right eye, with no attempt to balance the distribution of TMFs relative to the vertical meridian (a point discussed below).

### Apparatus

The stimuli, displayed on a conventional monitor (Dell 2407WFPHC, 1024 x 768 x 8 bits refreshed at 60 Hz) placed at 57 cm from the eyes, were generated by custom software (Jeda, Lorenceau & Humbert, 1990), running under Windows 10 (Microsoft Ldt, USA). Monocular eye-movements and pupil size were recorded at 500Hz with a Live Track Lightning remote eye-tracker (Cambridge Research System Ltd, https://www.crsltd.com/tools-for-vision-science/eye-tracking/livetrack-lightning/) placed at 30 cm from the eyes. Recordings were down sampled to 60 Hz for further analyzes. A chinrest was used to stabilize the participants’ eyes relative to the eye-tracker.

### Procedure

The session started with positioning the participants and checking the quality of the eye-tracker signals. A short 5 point calibration was performed once for the whole session. Note that precise calibration is not required, as the measure of pupil size is independent of calibration quality, and that relative eye-movements are sufficient to analyze fixation (in)stability. A short questionnaire was given to the participants to assess their general state (fatigue, existence of treatments, etc.), during which they adapted to the dim ambient light of the testing room. A session comprised 3 tests of 1 minute each, presented to both eyes, resulting in 6 minutes of recordings. One test was the mPFT grey test described herein. A second test used chromatic mPTF. The last test assessed the Pupil Cycle Time (PCT) using a computerized setting (See Ajasse et al. 2022). We here report the results for the grey mPFT test only. For all these tests, participants were simply asked to maintain fixation at the center of the screen, marked by a small colored circular fixation disk (0.2°), and to limit as much as possible the number of blinks, with no other concurrent task (the fixation disk randomly changed color –red, green, blue, yellow-every 2 seconds, allowing for the addition of a task, if needed, such as counting the number of occurrence of a specific color). A brief rest between the different runs was used to change the stimulated eye (Right or Left in alternation). We evaluated test-retest variability in some participants (N=8) by repeating the mPFT test 2 or 3 times on different days. Overall, a session lasted about 15 minutes (note that running the grey mPFT stimulus for both eyes takes only 2-3 minutes).

A mPFT run started with the onset of a centered fixation target, followed after 1 second by a brief full screen flash (166 msec.) at the maximum screen luminance (165 cd/m²) to elicit a pupil light reflex (PLR), followed by a dark screen for 3 seconds. Afterward, the static mPFT homogeneous grey stimulus (51 cd/m²) appeared on the screen for 3 seconds, allowing the pupil to adapt to the mean luminance, followed by 45 seconds of temporal modulations of the 9 sectors simultaneously.

### Participants

36 students (age range 20-29, 18 females) in the school of optometry of Université Paris-Saclay were enrolled in this study. All performed the pupillary tests described above as well as an Optical Coherence Tomography (OCT) and an ophthalmic examination as part of their university course. All participants gave informed written consent in accordance with the Declaration of Helsinki.

### Data analyzes

One data set from one participant was corrupted because of a technical issue and was removed from further analyzes. All remaining data were included in analyzes performed with Matlab R2018b (The MathWorks, Natick, MA, USA). The raw pupil data, down-sampled at 60 Hz, were first corrected for blinks and recordings artifacts. Blinks were detected as zeros in the pupillary traces. The corresponding blink data were replaced by a smoothed linear interpolation, using a pre- and post-blink offset of 4 samples (66.6 msec.). As the pupil is unlikely to change rapidly, we then performed a second pass to detect “artifactual transients” for which the pupillary signal was higher than a threshold, calculated using an *ad hoc* formula combining velocity and acceleration values, set once for the whole dataset. These “transient” data were replaced by a smoothed interpolation to avoid that they significantly impact the power spectrum computed afterward using the *fft* Matlab function.

As the initial and final pupillary responses were sometimes noisy, and to allow for pupil entrainment at the beginning of a run, the beginning and end of the pupillary signal was trimmed by 60 samples (1 sec.), so that 43 seconds of pupillary responses were used in analyses (2580 samples). The corrected pupillary signals were then z-scored, a ramping window of 100 samples, ranging from 0 to 1, was convolved with the beginning of the trace, and a window ranging from 1 to 0 was convolved with the end of the trace to avoid that abrupt onset and offset introduce spurious power in the Fast Fourier transform (FFT) that was computed on the so-corrected pupillary signals. Fourier analyzes were computed separately for each FOI. As a matter of fact, obtaining reliable amplitude and phase estimation for each FOI requires that the FOI to be analyzed is an integral multiple of the frequency resolution, FR (equal to the sampling frequency, SF, divided by the number of samples, N). This avoids that FOI power spreads across different bins of the spectrum. To that aim, we adjusted the frequency resolution, FR, by decreasing the number of samples, N, until this constraint was met: FR=SF/(N-x), with x adjusted separately for each FOI so that FOI/FR is an integral value (or very close to it, using a tolerance threshold equal to 10^-6^). Tests performed on the stimulus signal proved this method was efficient and reliable to precisely recover both the signal powers and phases. The amplitudes and phases of the pupillary signal obtained after adjusting the FR for each FOI in this way were stored. The phases were first computed for each FOI of the luminance modulations of the stimulus itself, so as to take the temporal offset (60 samples) into consideration. We then computed the relative phases between the stimulus and the pupillary signals for each FOI. The differences between the pupillary phases and the stimulus phases -phase shift at each FOI- were computed with the CircStat toolbox (Berens, 2009), and transformed into delays. In addition, the cross-correlation between the average time varying luminance modulation of the stimulus and the pupillary oscillations (after detrending the pupillary signal to suppress low frequency and linear drifts components) was computed to derive the lag between both signals, thus providing an overall estimate of the time needed for retinal processing, propagation time through the optic nerve and relay nuclei, as well as the constriction time constant of the iris sphincter and dilator muscles.

We observed that the spectrum of the pupillary signal tends to follow a 1/f distribution (Supplementary Figure 1A) and further contains “noisy” components, possibly related to imperfect signal correction at blinks. We therefore decided to normalize the raw spectral powers computed for each FOI. We did so by dividing the mean of the spectral power at bins surrounding the FOI 1/3x(SP(foi-1)+ SP(foi)+ SP(foi+1)) by the mean of the spectral power in surrounding bins 1/4x(SP(foi-3)+ SP(foi-2)+ SP(foi+2)+ SP(foi+3)). Normalizing the power spectra in this way takes into account the spurious noisy power that can exist for each frequency band, while homogenizing the power distribution of FOIs. Several other normalization procedures (by frequency or by regional bins of the spectrum) were tested, but appeared to be distort or skew the FOI power distribution for some participants (Supplementary Figure 1B,C,D).

We ascertain that peaks in individual spectral power were triggered by, and phase locked to, the stimulus’ FOIs, by comparing the spectral power of the pupillary responses averaged across all participants to the average of all individual power spectra. If individual pupillary responses were not phase locked to FOIs, random phase shifts would flattened the mean pupillary response, whose power spectrum would lack peaks at FOIs, or at least would decrease their respective power. This was not the case (Supplementary Figure 2), demonstrating that our stimulus did tag the pupillary response at each FOI in each case, although with varying power, depending on the region at stake (see Results). Time Frequency maps were also computed (with m=96) to verify that the power spectrum at FOIs was sustained during a run and did not result from short successive episodes of oscillatory activity at different times for different FOIs (Supplementary Figure 3).

Test-retest variability was assessed by computing the Pearson’s coefficient correlation between the FOI spectral power collected during different mPFT runs, as well as with Bland-Altman plots (Bland and Altman, 1983).

We also computed the correlations between the averaged retinal nerve fiber layer (RNFL) measured with OCT and the averaged spectral power, to evaluate whether the functional pupillary responses are related to the structural characteristics of the participants’ retinae.

## Results

All, but one, recordings of the 36 participants were included in analyzes (70 eyes), despite some recordings having a large number of blinks and data corrected for transients whose distributions are shown in Supplementary Figure 4 for all participants.

Figure 2 presents an example of the different steps of our pipeline analyzes that includes: **A**. Visual inspection of raw eye-movements, pupillary activity and technical event tracks. **B**. Analysis of the PLR, after blink detection and correction, from which 5 descriptive variables are derived (see Table 1 for the list of all features derived from analyzes). **C**. Analysis of eye-movements –fixation (in)stability-during the stimulation, from which 6 variables are computed. **D**. **1.** Delineation of the relevant pupillary signal during mPFT stimulation, correction of blinks and transient data, with computation of 7 descriptive variables, and characterization of 5 global pupillary variables extracted from the corrected pupillary signals including the stimulus/signal cross-correlation lag. **2**. FFT analyzes, estimating the amplitude and **3** phase of each FOI (black symbols: 9 stimulus phases, 9 pupillary phases, open blue symbols: 9 Periods, filled blue symbols: 9 stimulus/pupillary phase differences). **E**. Maps of the Pupillary Response Fields for the raw and normalized amplitude as well as phase lag, according to the sectors of the mPFT stimulation. As it can be seen in Figure 2D, the pupillary spectral power (PSP) of the sustained pupillary response shows peaks at each of the 9 FOIs, indicating that the pupillary response was modulated by each sector.

**Figure 2:**
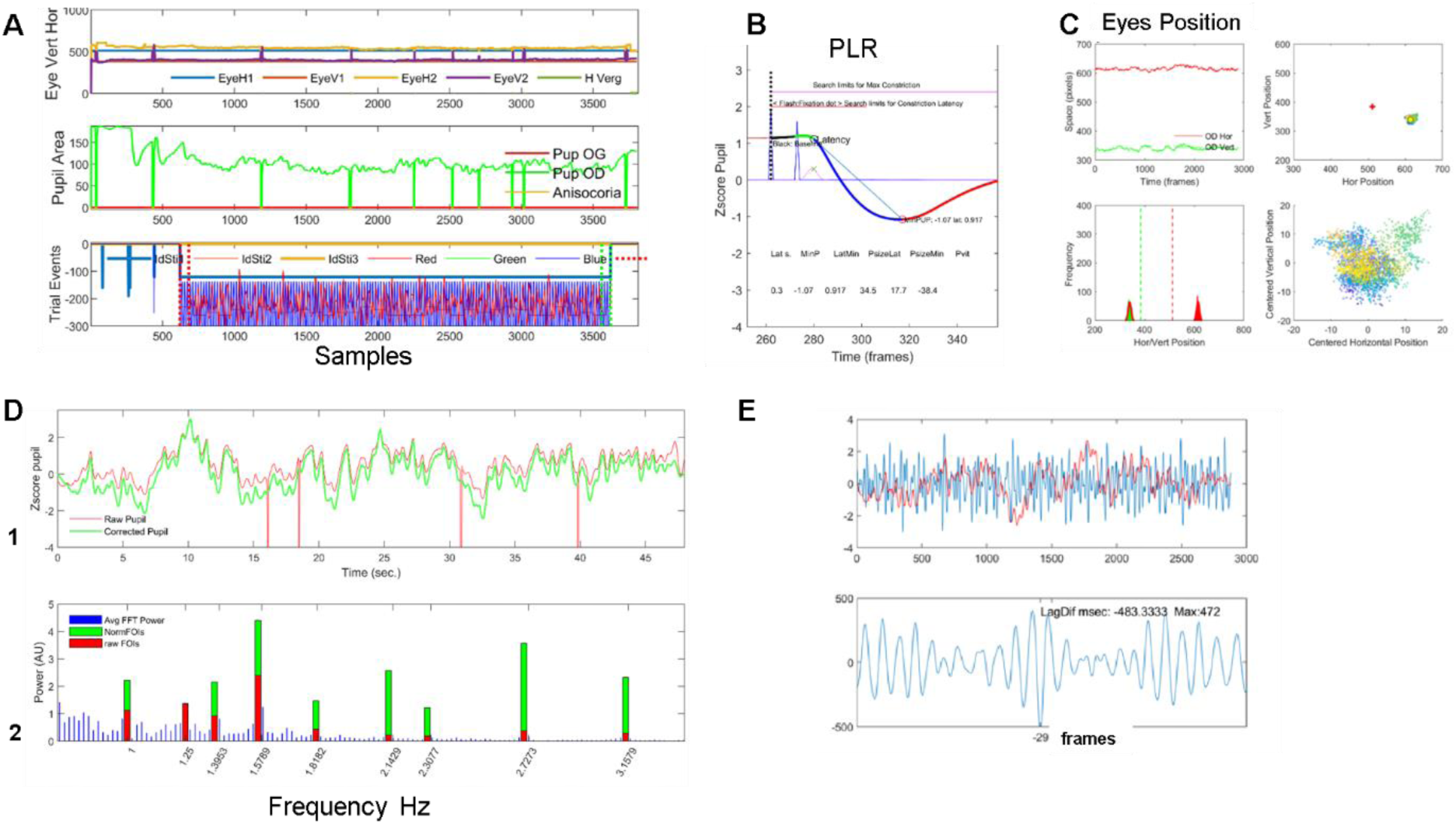
Steps of analyzes: **A**. Visual inspection of raw eye-movements, pupillary activity and technical event tracks. **B**. Analysis of the PLR, after blink detection and correction, from which 5 descriptive variables are derived (see Table 1 for the list of all features derived from analyzes). **C**. Analysis of eye-movements –fixation (in)stability-during the stimulation, from which 6 variables are computed. Top left: Position over time; Top right: Position over space, whole screen; Bottom left: histogram of vertical and horizontal eye positions.Bottom right: zoom on centered spatial eye positions **D**. **1.** Raw (red trace) and corrected pupillary signal (green trace) during mPFT stimulation, correction of blinks and spurious data, with computation of 7 descriptive variables, and characterization of 5 global pupillary variables extracted from the corrected pupillary signals including the stimulus/signal cross-correlation lag. **2**. FFT analyzes, estimating the amplitude spectrum: mean FFT (thin blue lines); Raw FOI power (red bars); Normalized FOI power (Green bars) **E**. Cross-correlation between the luminance oscillations of the stimulus and the pupillary response. Top Stimulus oscillation (blue line) and pupillary response (red line). Bottom, cross-correlogram results.

**Table 1:**
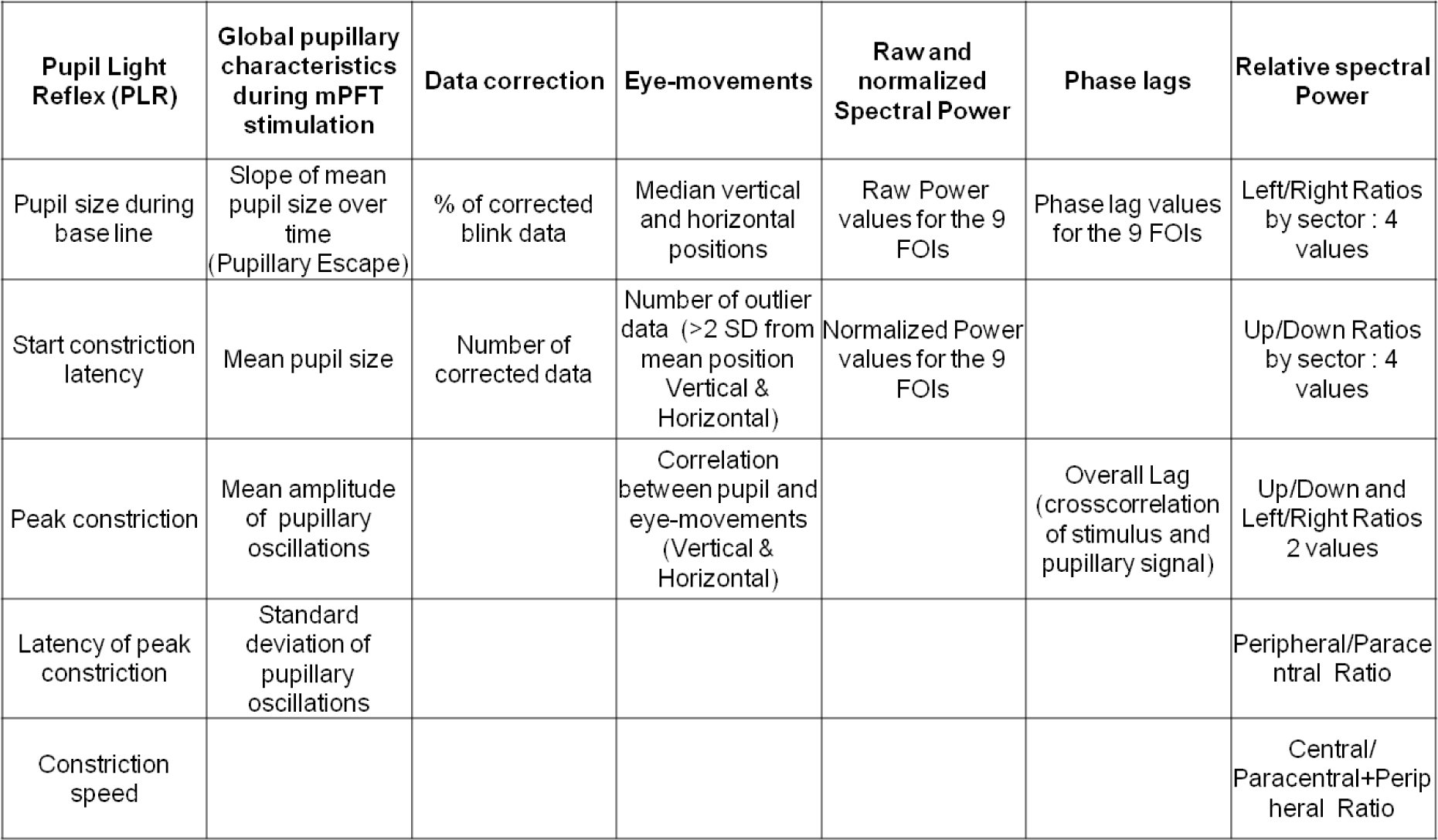
List of the variables derived from the analysis of each run.

For each run, we computed 55 variables characterizing pupil and eye-movement activity during the mPFT stimulation (Table 1).

Figure 4 show the group results for the right and left eye. Figure 4A shows the correspondence between the Stimulus Sectors and the associated FOIs (Hz). Figure 4B shows the distribution of raw power for the 9 FOIs and associated PRFs. Figure 4C shows the distribution of normalized power for the 9 FOIs and associated PRFs. Figure 4D shows the distribution of phase lags for the 9 FOIs and associated PRFs. In this later graph, the large variability observed for 2 FOIs reflects the presence of aberrant values for some participants, again reflecting the circular nature of phase computation with FFT.

### Distribution of Pupillary Spectral Power

The power distribution and Pupillary Response Fields (PRF) in Figure 4 show that the pupillary spectral power (PSP) is heterogeneous across sectors –hence across the visual field-, with a larger raw power for peripheral sectors than for paracentral and central sectors, and, in particular, for the sector projecting onto the infero-nasal retinae. The distribution of PSPs and phase lags is however similar across individuals, indicating that Pupillary Response Fields computed with our method reflect a common spatio-temporal organization of retino-pupillary circuits, despite some idiosyncrasies. This homogeneity at the group level also suggests that short episodes of fixational instability or blinks that differ amongst individuals did not significantly perturb the sustained pupillary response during a run (see discussion of this aspect below).

One possible account of the PSP differences across sectors could be related to the different TMFs used for each sector. This seems unlikely for several reasons. First, the differences in TMFs being small –from 1 to 1.58 Hz for the peripheral sectors-, it is unlikely that the pupil temporal frequency transfer function changes much over such a small range (see Clarke et al., 2003). Second, the fact that the power distribution differs in the right and left eyes despite the TMFs being similarly distributed argues against a pure contribution of TMFs. Third, it can be seen on Figure 3 that the largest power is found for 1.58 Hz for the right eye, similar to that obtained at 1.58 Hz for the left eye. Unless one postulates a very strange temporal transfer function with ups and downs, the data suggest that temporal frequency *per se* does not explain the power distribution across sectors.

**Figure 3:**
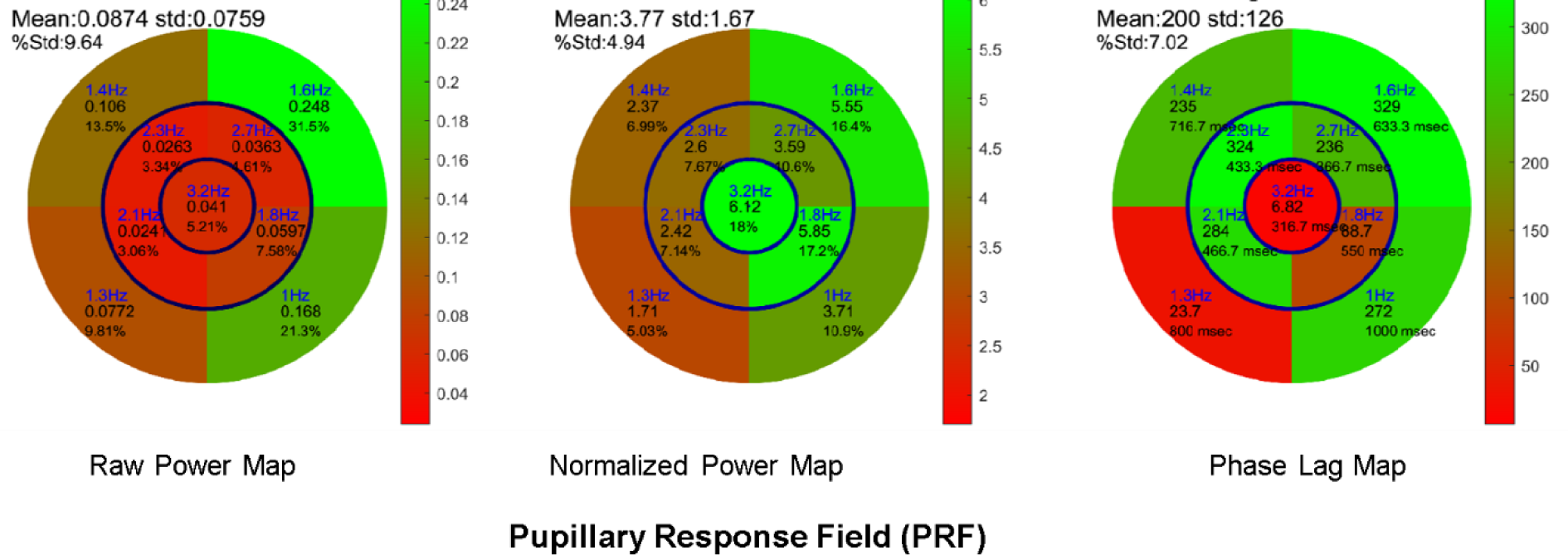
Maps of an individual Pupillary Response Fields for the raw power (left), normalized power (middle), and phase lag (right), according to the sectors of the mPFT stimulation Numbers in each sector indicate the TMF, FOI power and the percentage of pupillary activity for that sector, relative to the sum of all FOI power. For the phase map, the numbers in each sector indicate the TMF, the phase lag in degrees, and the corresponding latency in msec.

Rather, this spatial distribution likely reflects the intrinsic pupillary sensitivity of different retinal regions, in line with other studies showing that the infero-nasal retina, on which the upper-temporal sector projects, entrains larger pupillary responses than other, -non-foveal-regions (Wilhelm et al., 2000; Hong et al., 2001; Maddess et al., 2009). This finding is also in line with the non-uniform distribution of human ipRGCs characterized by a larger density in the infero-nasal retina (Hannibal, et al., 2017).

One exception is the relatively small raw power observed for the central disk sector. It is known that light stimulation of the macula elicit large pupillary responses, as compared to paracentral and peripheral stimulations (Hong et al., 2001). This large pupillary response to central stimulation is further corroborated by a dense distribution of ipRGCs around the macula (Hannibal, et al., 2017). Here, the central region is coupled with the highest TMF, 3.17 Hz, for which the temporal transfer function of the pupil starts to decline (Clarke et al., 2003). As a matter of fact, our choice of coupling the highest TMF to the central region was dictated by this decreased response to “high” temporal frequency, so as to limit the contribution of macular stimulation to the overall pupillary activity, which could otherwise limit or dominate the contribution of other regions.

### Timing of Pupillary Oscillations

Of interest is the distribution of phase lags presented in Figure 4, as it could reveal retino-pupillary processing delays, including retinal temporal integration, propagation times through the optic nerve and nuclei lying between the retina and the iris muscles (see Szabadi and Bradshaw, 1996), as well as the mechanistic time constant of iris motility. Although the phase lags shown here are very different across sectors, they are very similar across participants, with few exceptions (mostly related to the circular nature of phase computation). The lag distributions for the right and left eye are similar, indicating phase lags are related to the TMFs, independently of the sector locations. These distributions are irregular (in terms of phase and delay), and are presumably related to the circular nature of phase processing with FFT. Correcting these values to match the hypothesis that too short or too long phase lags are unlikely would require several assumptions regarding the expected timing of pupillary responses. We preferred not to consider that the computed phase lags reflect veridical physiological lags, as these values are at odd with reports of PLR latencies, and are sometimes positive which is very unlikely, but may still be relevant for assessing differences between healthy individuals and patients in clinical studies (see below). In our view, the phase lags computed herein just indicate that the mPFT stimulus did entrain pupillary oscillations in a similar way for all participants.

**Figure 4:**
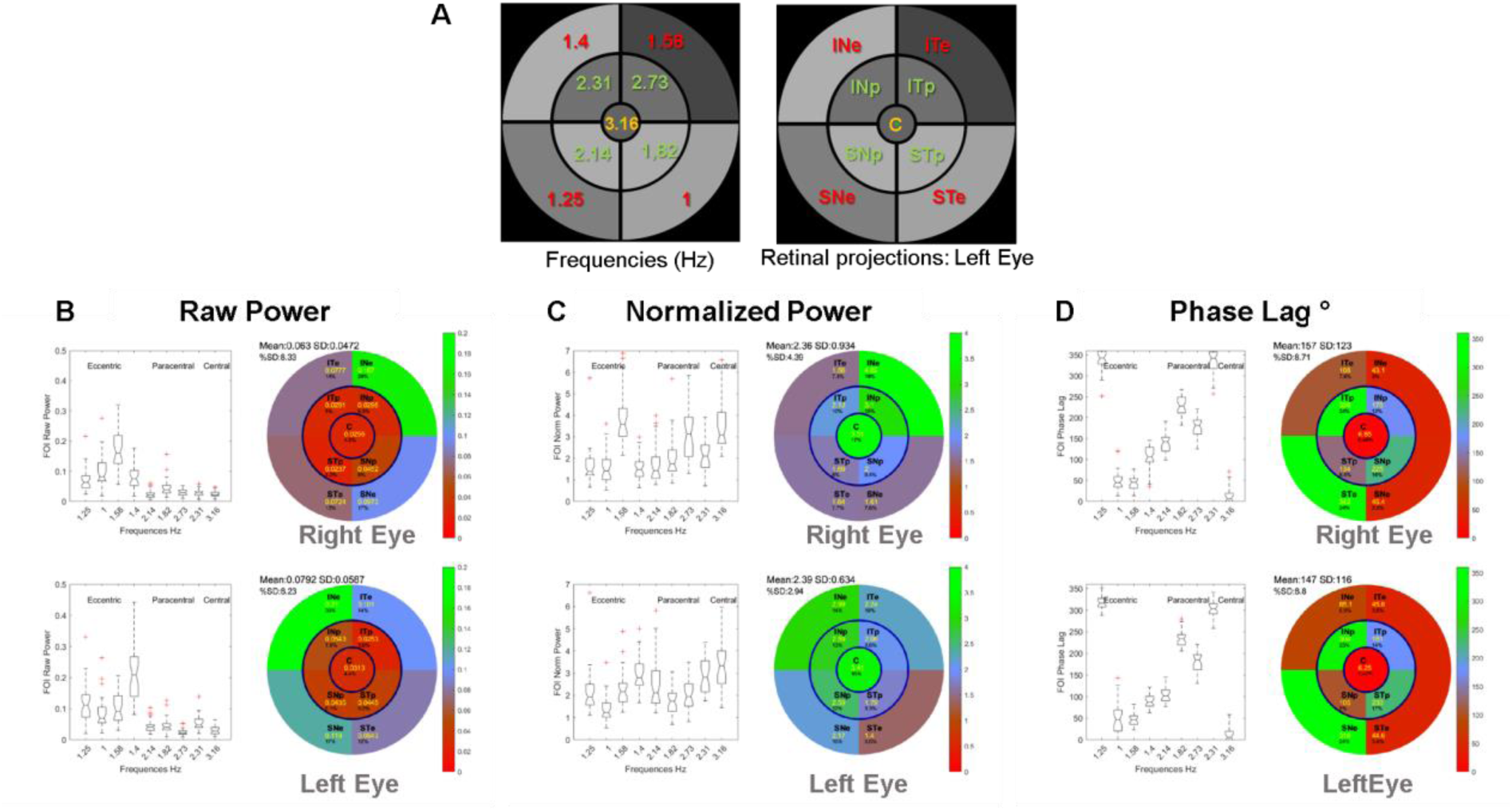
Group results showing the power distribution and Pupillary Response Fields (PRF) of the right and Left Eyes: **A**: Stimulus Sectors, associated FOIs (Hz) and retinal projections. **B**: Distribution of raw power for the 9 FOIs and associated PRFs. **C**: Distribution of normalized power for the 9 FOIs and associated PRFs. **D**: Distribution of phase lags for the 9 FOIs and associated PRFs.

Computing stimulus/signal cross-correlation seems a better way of assessing the overall latency of the pupillary responses relative to the mPFT stimulus. These measures give values around 500 msec. (range 420-540 msec.) with little inter-individual variability (Figure 5), and are comparable to the maximum constriction latencies reported elsewhere for the PLR, ranging between 500 and 600 msec (Ellis, 1981; Link and Stark, 1988; Whilem et al. 2000; Tan et al., 2001).

**Figure 5:**
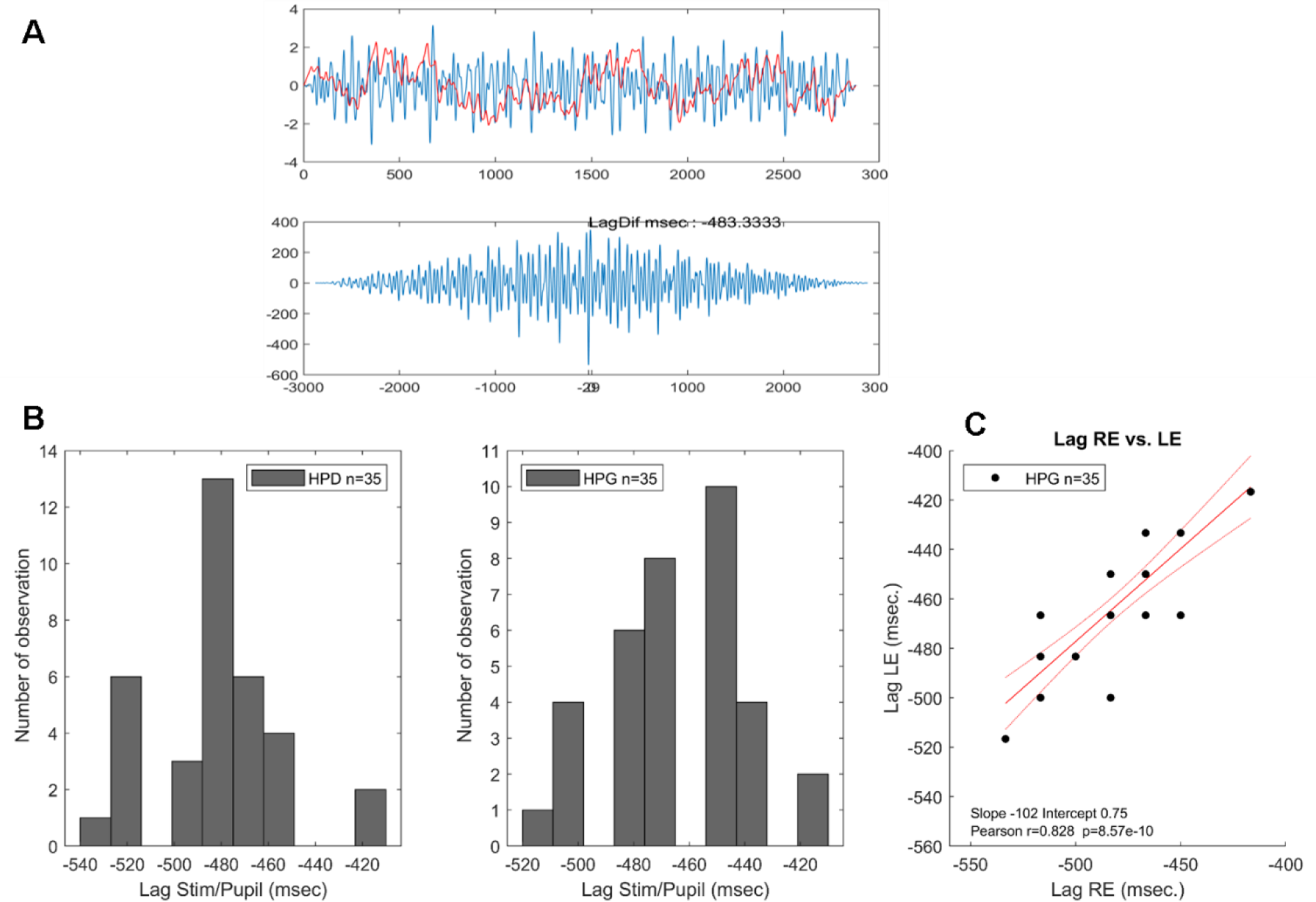
**A:** Example of a Stimulus/Pupillary signal cross-correlation using the Matlab xcor function. **B:** Histogram of phase lags for all participants. Left: Right Eye; Middle: Left Eye. **C:** Correlation between phase lags of the Right and Left eyes. Note that because pupillary signals are down sampled to 60 Hz, the time resolution of lags is only 16.666 msec.

The extent to which the latencies of the PLR recorded at the beginning of a run (see Method and Table 1) and the estimates derived from the Stimulus/Signal cross-correlation are correlated is shown in Figure 6.

**Figure 6:**
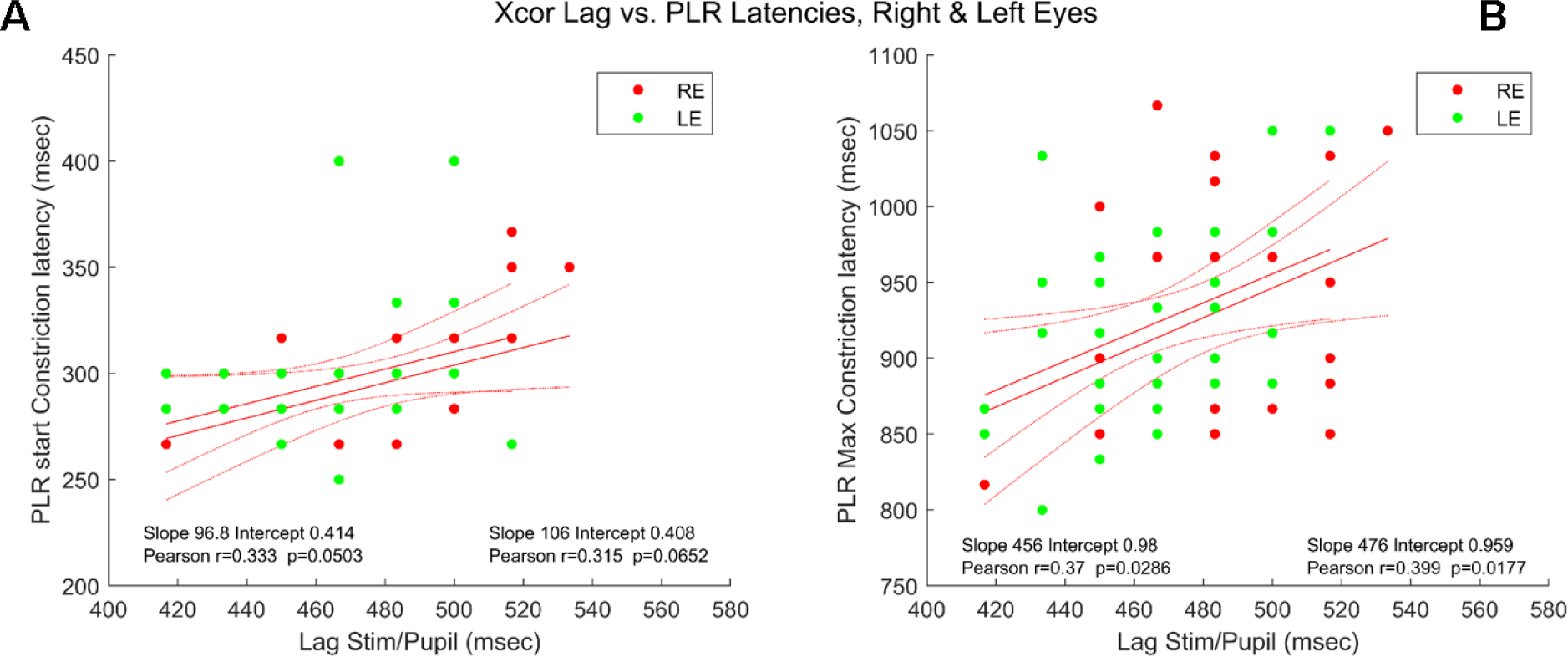
Correlations between the Stimulus/Signal lag and PLR latencies. **A**. Lag vs. PLR start constriction latency: Right (red disks) and Left (green disks) Eyes. **B**. Lag vs. PLR maximum constriction latency: Right (red disks) and Left (green disks) Eyes. Inserts indicate the values of Pearson coefficient correlation for each Eye.

Figure 6A shows the latency of the beginning of pupil constriction elicited by a flash as a function of the lag during sustained stimulation; Figure 6B shows the latency of the constriction peak as a function of the lag during sustained stimulation.

As it can be seen in Figure 6, the better –and significant – correlations are obtained between the Stimulus/Pupil lag and the latency of the maximum PLR constriction for both eyes. This suggests that the measure of Stimulus/Pupil lag, as done here, does reflect a relevant aspect of the timing of pupillary responses during sustained stimulation, capturing some of the inter-individual differences found with the PLR. Note that the average Stimulus/Signal lag (482.4 and 463.3 msec. for the right and left eyes, respectively) are shorter, but still in the range of the PLR latencies reported in the literature (about 550 msec in Wilhelm et al. 2000, or Tan et al., 2001), despite very different stimulation conditions.

Note that in studies using the PLR, the latency of maximum constriction is computed after dark adaptation, such that pupil baseline size is at its maximum. In contrast, the lag computed with a sustained mPFT stimulus reflects the continuous adaptation of pupil size to varying luminance intensities, mixing dilation and constriction, which could account for the differences reported here, although direct comparisons of PLR latencies with the lags found with mPFT should be considered with caution.

### Test / Retest with mPFT

Determining whether mPFT is reliable and stable is important if it were to be routinely used for clinical assessments. To evaluate the repeatability of mPFT, several participants (N=8) repeated the protocol 2 times on different days and at different times. We then computed the correlations of the pupillary features extracted during data analyzes for each participant and at the group level. Correlations and Bland-Altman plots between Run1 and Run2 were computed for the Raw Power, the Normalized Power, the PLR and global Pupillary variables (see Table 1 for a detailed list).

Figure 7 shows examples of correlations for 2 individuals and different pupillary variables. The test/restest correlation coefficient for the raw power of the 9 FOIs is high (>0.8, (Fig 7a) and larger than that found for the Normalized Power. Figure 8 shows the distribution of Pearson’s correlation coefficients for each of the 8 participants for different variables (Raw and normalized power, Phase lags and PLR). With some exceptions for the Normalized power and phase lags, test/retest correlations are high for all participants. Finally, Figure 9 shows correlations of the Raw (r=0.82) and Normalized Power (r=0.71) at FOIs and Bland-Altman plots at the group level for the Raw and Normalized Power.

**Figure 7:**
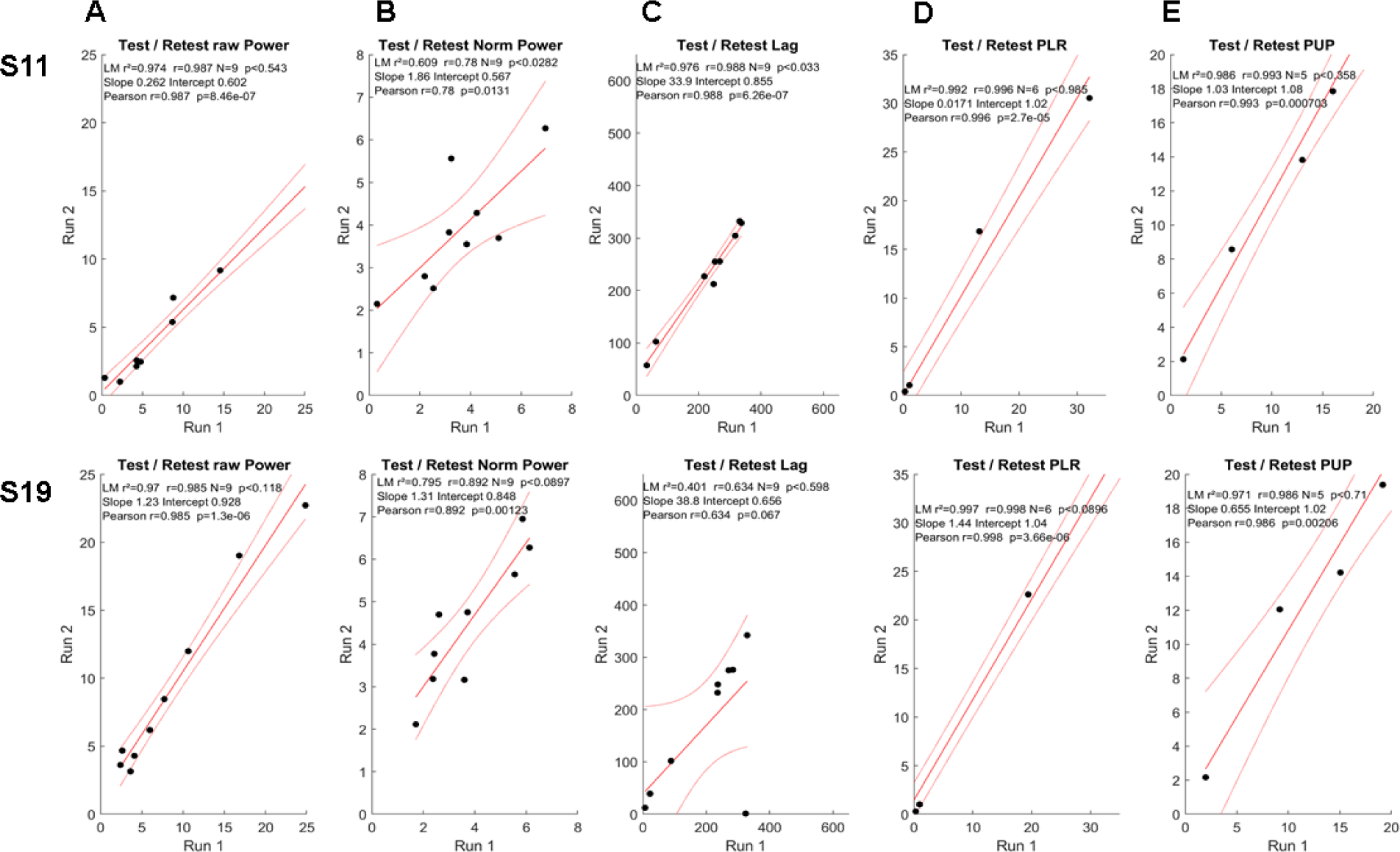
Examples of test/retest data for 2 participants. **A**. Correlations for the Raw Power; **B**. Correlations for the Normalized Power. **C**. Correlations for the Phase Lags. **D**. Correlations for the PLR variables. **E.** Correlations for the Global pupillary variables. Inserts indicate the values of Pearson coefficient correlation for each Eye.

**Figure 8:**
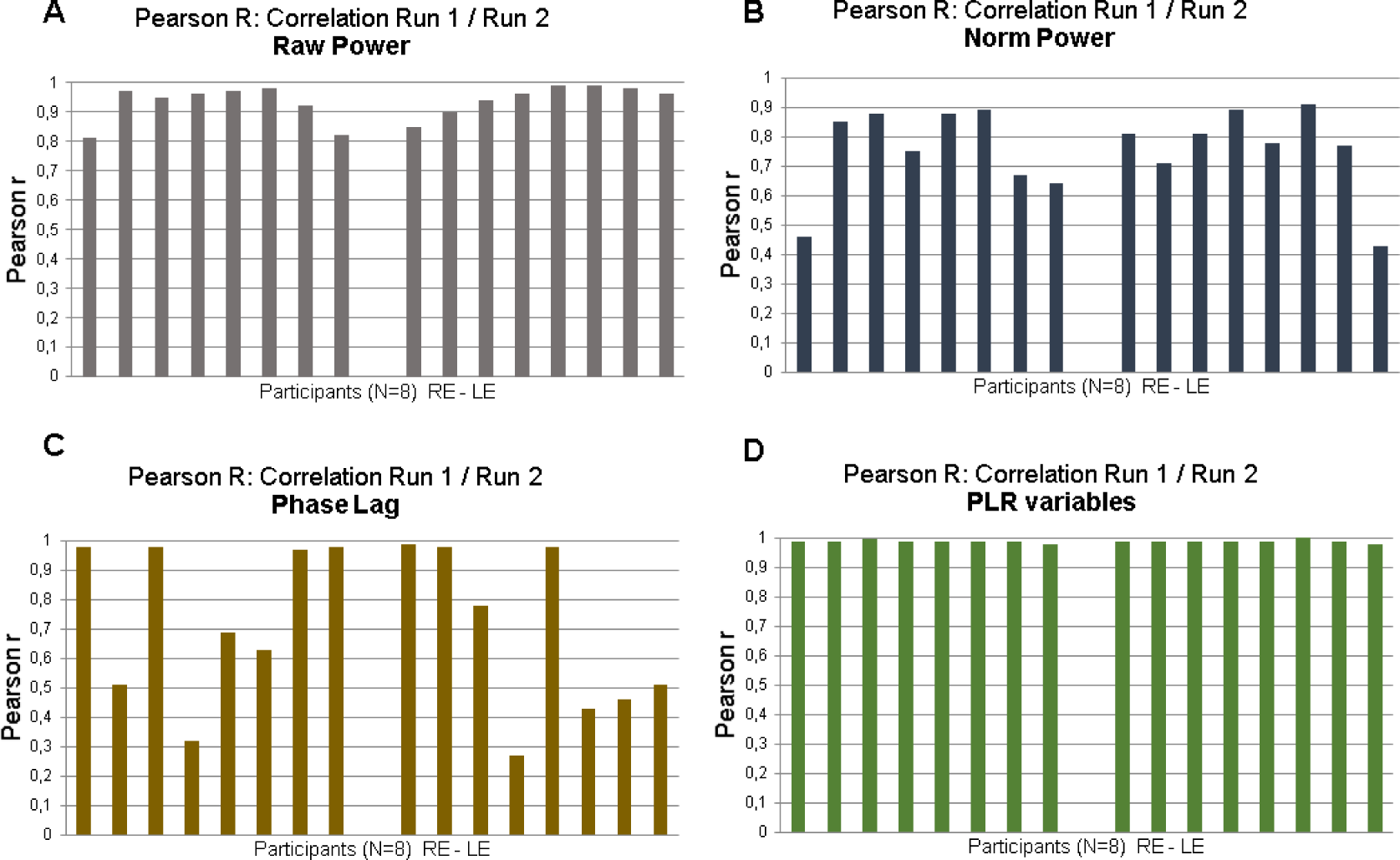
Distribution of Pearson’s r coefficient of 8 participants for the Right and Left Eyes. **A**. Correlations for the Raw Power; **B**. Correlations for the Normalized Power. **C**. Correlations for the Phase Lags. **D**. Correlations for the PLR variables.

**Figure 9:**
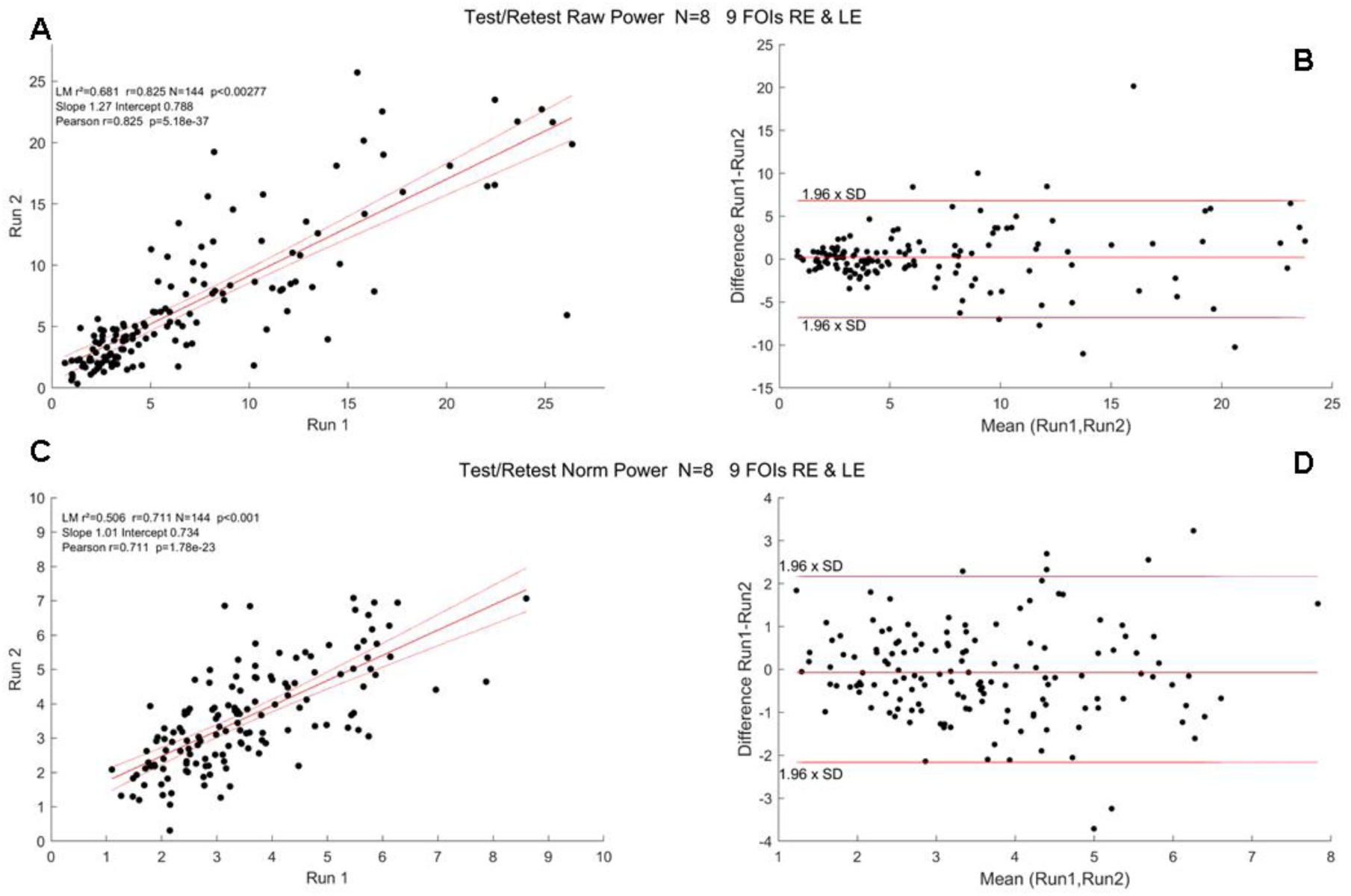
Test/Retest Pearson’s r coefficient at the group level (N=8, pooled Right and Left Eyes). **A**. Correlations for the Raw Power; **B**. Bland-Altman plot the RawPower. **C**. Correlations for the Normalized Power. **D**. Bland-Altman plot the Normalized Power.

These test/retest analyzes (with a delay of at least 1 day between Run 1 and Run 2) all indicates that the variables derived from mPFT are stable over time. It can be noted, however, that correlations are better for the Raw, as compared to the Normalized, FOIs Power, presumably because the power normalization done here takes into account the residual power of the FFTs (see Data Analyzes) that can vary depending of the corrections of blinks and transients that introduce small artifacts that introduce power at all frequencies. Correlations values are smaller for the phase lags. This is presumably due to the fact that the estimates of some phase lags are biased by the circular nature of this variable (see above Timing of mPFT).

### Correlations between Spectral Power and RNFL thickness in young healthy participants

Several clinical studies report correlations between pupillary features of the PLR and Retinal Nerve Fiber Layer (RNFL) estimated with Optical Coherence Tomography (Chang et al., 2013; Najar et al., 2018; Gracitelli et al., 2014; Rukmini et al., 2019; Stelandre et al., 2023; Ajasse et al., 2022). In these studies, depending on the protocol and stimuli at stake, different features of pupillary signals from populations of patients with ophthalmic pathologies are measured and compared to structural characteristics of the retina. To our knowledge, no such correlation has been looked for in a cohort of young healthy participants to evaluate the existence of functional/structural correlations. In order to perform such comparison, we first decoded the RNFL values from the PDF files that summarize the OCT results (Spectralis, Heidelberg Engineering, Germany) of each participant and plotted the averaged RNFL values as a function of the averaged spectral power (Figure 10).

**Figure 10:**
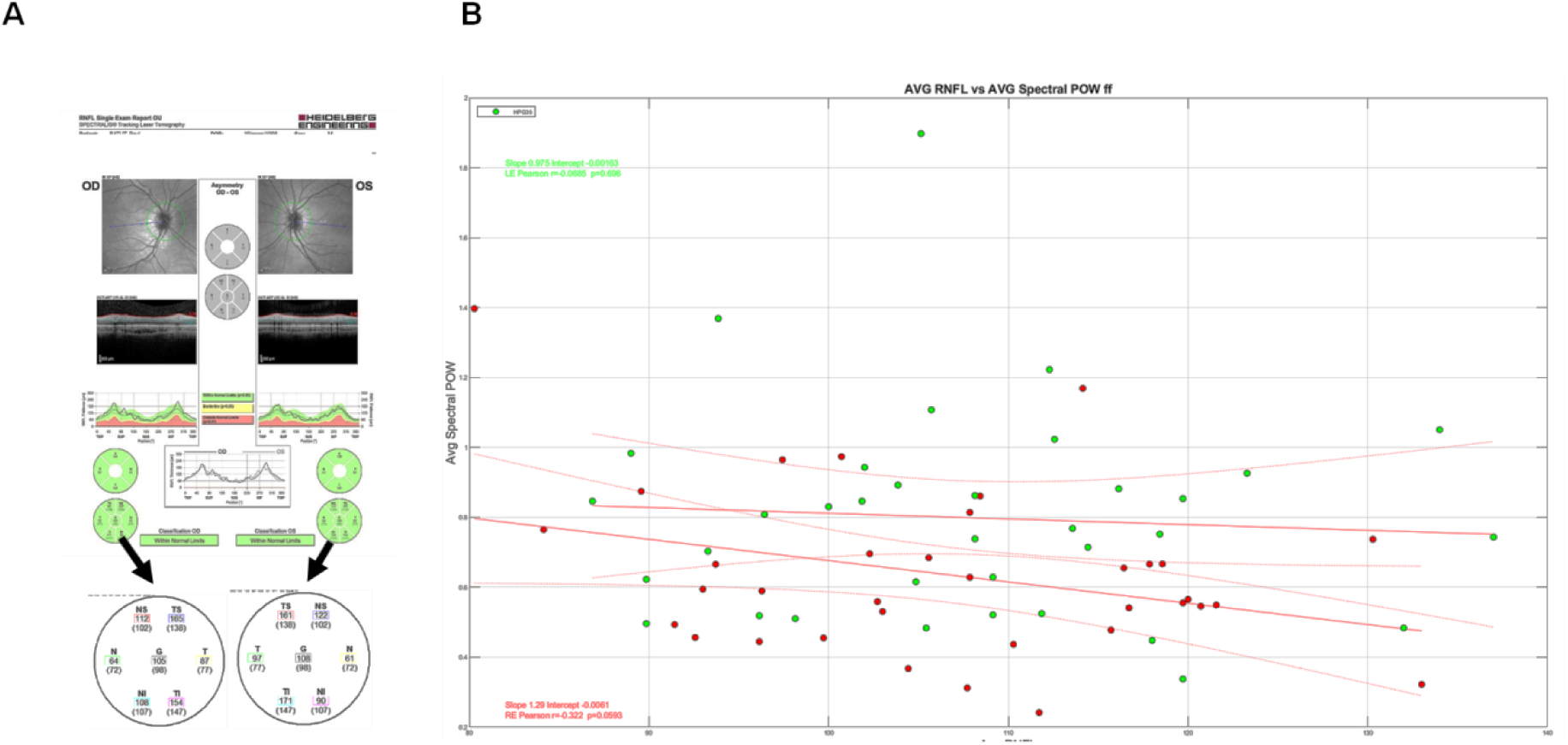
**A**. Example of RNFL data and extraction of relevant values from a PDF file. **B**. Distribution of the average mPFT raw power as a function of the average RNFL values for the right eye (Red symbols) and the left eye (Green symbols). No correlation is found between both variables (Inserts are the Pearson’s correlation coefficients, with different colors for the 2 eyes).

As it can be seen in Figure 10, the averaged RNFL varies from about 80 µm to more than 130 µm in this population of young healthy participants. Similarly, the averaged mPFT powers are different across individuals. However, we do not observe a significant correlation between the averaged RNFL and averaged mPFT power. This suggests either that the thickness of the RNFL is not important for the functioning of the pupillary circuits, at least in the observed range, or the fact that comparing the average values of RNFL and mPFT power is irrelevant because the means do not capture the intra-individual variability of each distribution. We attempted to refine this analysis by comparing the mean RNFL values of the upper retina (TS, NS, see Figure 10A) with the mPFT power of the lower sectors, and the lower RNFL retinae values (TI, NI) with the mPFT power of the upper sectors. These comparisons did not reveal significant correlations of both variables, possibly because the location of these measures does not tightly coincide.

So, contrary to clinical studies conducted on some pathologies (Glaucoma in particular) showing correlations between RNFL and pupillary responses (Chang et al., 2013; Najar et al., 2018; Rukmini et al., 2019; Stelandre et al., 2023), the same does not hold for young healthy subjects. We hypothesize that RNFL/Pupillary responses correlations can be found only for extreme values (very thin RNFL for advanced pathologies), but not in healthy subjects.

### No effects of gender on mPFT power

Anisocoria, the physiological difference between the size of right and left pupils (static anisocoria) or between the pupillary responses of the right and left eyes to light stimulation (dynamic anisocoria), may differ between male and female (for dynamic anisocoria in particular) in 20 to 30% of the population, with a prevalence increasing with age (Poynter, 2019). To determine whether sex could influence the responses to mPFT, we compared the distribution of spectral power of the right and left eyes in our population (18 females). Although we found differences between the two eyes, these were not related to gender (p>0.05).

## Discussion

We now discuss the results presented above, and the benefits as well as limitations of our approach, before summarizing the results of some clinical studies performed using mPFT.

The mPFT method presented here allows objectively assessing multifocal Pupillary Response Fields reflecting the integrity and functioning of retino-pupillary circuits in a short amount of time (1 minute per eye). This method permits to analyze both the power and phase lag for each FOI, and the overall latency of sustained pupillary responses to continuous luminance oscillations, together with several other variables, including global pupil size and reactivity, amount of blinks or fixational eye-movements stability. Test/retest done on different days on a subset of participants indicate that the distribution of mPFT power is stable over time, suggesting that mPFT results are robust.

Importantly, the Pupillary Response Fields permit to analyze the *relative* contribution of each sector to the overall pupillary response recorded at once, and not only the *absolute* power for each sector, allowing to identify sectors with relatively less power than others, an important feature in the perspective of using mPFT with patients presenting localized visual defects (scotoma), as looking for relative differences between sectors renders mPFT less prone to the influences of exogenous factors (e.g. ambient luminance, time of the day) and endogenous factors (medication or age for instance) that may modulate the mean pupillary response. Although we did not extensively present and analyze the distribution of *relative* spectral power herein (but see below for an example), we did compute 12 ratios between sectors or group of sectors that can be used in clinical studies, as described in supplementary Figure 5. As a matter of fact using this distribution of ratios did improve classification results in clinical studies.

The use of a continuous sustained stimulation also allows to evaluate additional variables collected within a single run, such as the number of blinks, the fixational instability and “spurious” eye-movements (saccades), known to vary with clinical conditions, and thus relevant to assess the existence of ophthalmologic issues.

### Effects of blinks and eye-movements and of the poor quality for some recordings

One may wonder whether the correction of blinks and transient data that are replaced by smoothed linear interpolations before the FFT computation (see Method), influences the overall power at FOIs. Plotting the percentage of Blink corrected data and number of transient corrected data as a function of the average spectral power (Figure 11) does not reveal any strong correlation, as some participants with numerous corrections have a larger averaged power that some participants with little or no corrected data.

**Figure 11:**
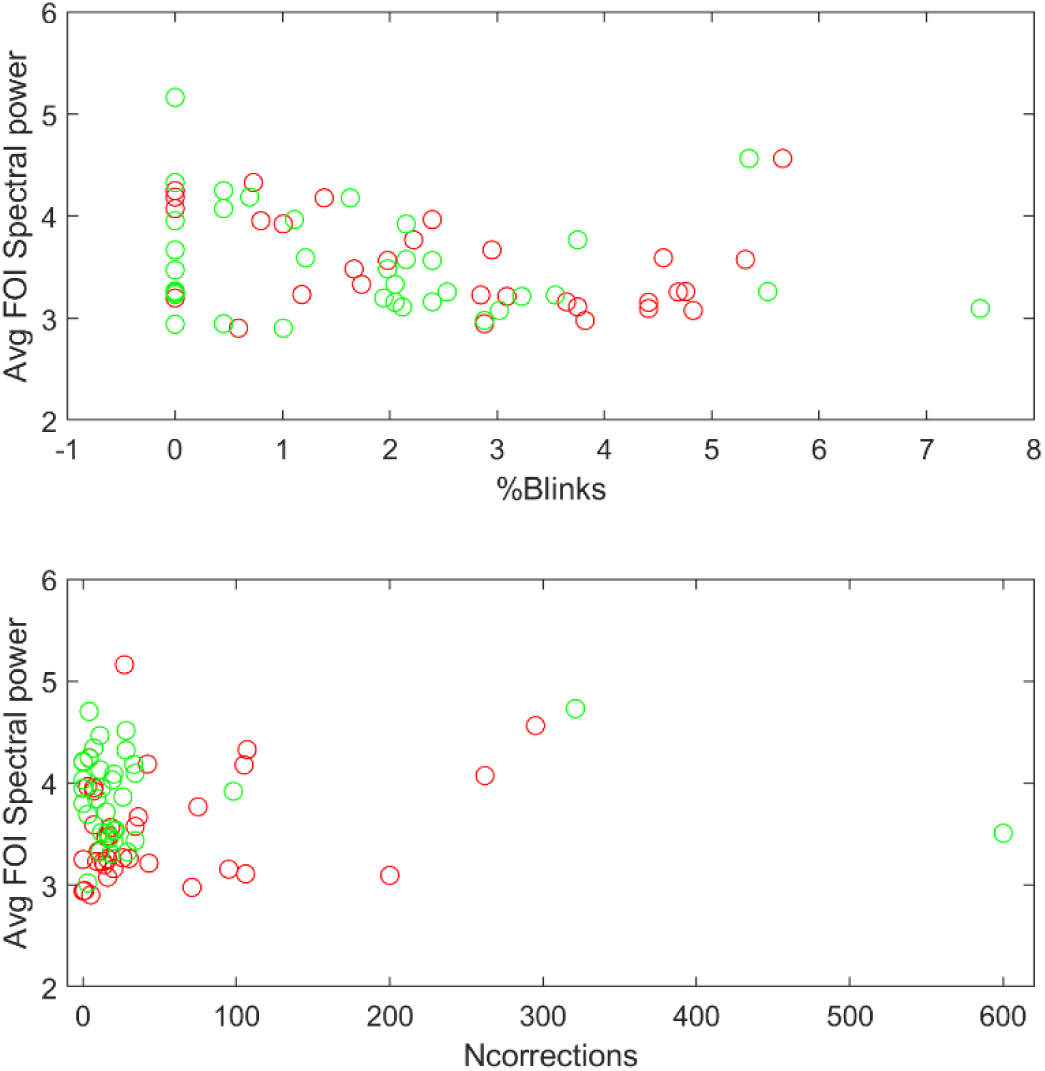
**A.** Averaged spectral power as a function of the percentage of blink corrected data. **B.** Averaged spectral power as a function of the number of data corrected for transients.

### Factors determining pupillary activity and choice of settings

Three factors may determine the amplitude of the spectral power at FOIs: the relative surface and orientation of each sector, the amplitude of the luminance modulations and the TMF associated to each sector.

Changing one or several of these attributes is expected to modify the relative or absolute contribution of each sector to the overall pupillary response. Although we tried to optimize these parameters for each attribute, other configurations are possible, and may be desirable, for instance to better homogenize the contributions of each sector. As a matter of fact, the distribution of raw FOIs’ powers (Figure 4) indicates that the large peripheral sectors tend to dominate the pupillary response. Changing the relative size of each sector, for instance reducing the size of the peripheral sectors relative the paracentral and central sectors, may better balance the distribution of power across sectors. Alternately, one could change the relative amplitude of the luminance modulations of peripheral, paracentral and central sectors, while keeping their sizes as they are. Note that the amplitude of luminance modulations should remain at a reasonable level to still elicit reliable and sustained pupillary activity.

Changing the TMF distribution, for instance by coupling a different temporal frequency with a given sector, does not deeply modify the spatial distribution of power, as was observed in control experiments (data not shown), strengthening the view that temporal frequency is not the determining factor accounting for the power distribution across FOIs, at least for TMFs below 4 Hz.

### Luminance versus RGB modulations of stimulus sectors

In this study, we continuously modulated the luminance intensities of each sector by changing the RGB values on each frame, and did not use luminance modulations expressed in cd/m² (or any other unit of light intensities). Thus, the luminance modulation relative to the mean luminance were imbalanced if expressed in cd/m² (see Method), because the luminance is exponentially related to the RGB units. Although we did not test mPFT with symmetrical, gamma corrected, luminance modulations –relative to the mean luminance-, we reasoned that the retina itself performing a logarithmic luminance compression, it may not be necessary –and possibly not desirable-to use a symmetrical luminance modulation. In addition, because the luminance modulations used here are of modest amplitude (from 67 to 187 in RGB units, corresponding to a 20 to 100 cd/m² luminance modulation relative to a mean luminance of 51 cd/m²) the resulting asymmetry also remains in a reasonable range. If the imbalanced luminance modulations had an impact of pupillary responses, one would expect to observe pupillary power at frequencies other than the FOIs. Finally, we tested mPFT with different screens having different settings (luminance, contrast or color temperature) with little impact on the relative distribution of power at FOIs. This indicates that mPFT is robust to changes in these settings, making it usable with standard hardware without the need of precise and constraining adjustments.

### Intermodulation products and pupillary responses

To determine whether reliable power exists at frequencies other than FOIs, we computed the average of all Fourier spectra from all participants (Supplementary Figure 6). Although the mean spectrum shows power at the 9 FOIs, significant power at other temporal frequencies also exist. In particular, for the right eye, the powers at 1.146 Hz, 0.667 and 0.4167 Hz are close to the power at FOIs. For the left eye, the powers at 0.667 and 0.4167 Hz are close to that for FOIs, and a small peak at 1.146 Hz is present. Other peaks of small amplitudes can be observed at different spectrum locations. It is unclear whether these peaks correspond to intermodulation products that would indicate a non-linear combination of 2 or more frequencies, or whether the power at these frequencies are unrelated to FOIs and results solely from frequency noise in the low frequency range. The fact that peaks are found at 0.4167, 0.667 and 1.146 Hz and in other similar locations in both eyes, although with different amplitudes, nevertheless suggests that power at these frequencies could be related to non-linearities in the pupillary response. Although it is challenging to characterize these non-linearities and to which FOIs are involved in producing these components, it could be interesting of analyzing them further, particularly in clinical conditions. Another possibility is that these low frequency peaks result from the imbalance of luminance osicllations relative to the mean luminance (see above)

### Effects of attention on mPFT spectral power

As mentioned in the introduction, focused attention and other cognitive factors induce a pupillary dilation, although of modest amplitude (Mathot, 2018; Hupé et al., 2012; Ajasse et al., 2017) through activation of the sympathetic pathway which in turn sends inhibitory projections onto the Edinger-Westphal nucleus driving the pupil (Szabadi & Bradshaw, 1996). Could attention modulate the spectral power of mPFT? Although attentional lapses or focused attention onto a sector could modulate pupillary activity, these pupillary modulations are unlikely to counterbalance the continuous strong and sustained visual drive of mPFT. For attention to significantly modulate pupillary activity may require sustained covert attention –to one or several sectors-during the whole test, which appears difficult to perform, and therefore unlikely. To nevertheless test for this possibility, we conducted experiments with young healthy participants (N=16, different from those who performed in the present study), whose task was to count the number of times two colored disks displayed side-by-side onto one sector had the same hue. The colors of the disks changed every second. Each of the 45 seconds trials (one for each sector, except the central disk, resulting in 8 trials) thus required covertly attending to a single location during the whole duration of the test, and involved counting and memory, all being cognitive tasks known to entrain a pupil dilation (see e.g. Mathot, 2018). A control condition used the same stimuli, but with no associated task (passive fixation). The experiment was done with 2 luminance modulation amplitudes (large, as used herein, vs. half the amplitude modulation used here) to determine whether covert attention is more likely to influence the spectral power when the visual drive is less strong. We did find a significant increase of the mean spectral power in the attention, relative to the no task, condition (Student T test, p<0.02), that was larger for a smaller visual drive (Cohen’d test: 0.75 for large luminance modulation, vs. 0.44 for the lower luminance modulation). However, we did not find significant selective increases of spectral power associated to the sector where the colored disk stimuli were displayed. As we used a visual stimulus –colored disks superimposed to the sectors-that could itself modulate pupil size, we performed an additional experiment using auditory signals, in which participants had to count the number of sounds with a higher pitch (460 Hz) than a reference (440 Hz, with 25% of sounds with a higher pitch). This experiment did not reveal any significant effect of the auditory task on the averaged mPFT power (Student T test, p>0.05, unpublished report, 2018).

Thus, sustained attention does modulate the average mPFT spectral power, but this effect is global and not spatially selective, such that attention does not appear to significantly modulate the distribution of *relative* power. Note that the two tasks were demanding and difficult to perform. It therefore seems unlikely that similar attentional effects would affect the spectral power when the task is simply to maintain fixation at the center of the mPFT stimulus, and the luminance modulation amplitude is large.

### mPFT and dark adaptation

In most studies using the PLR to assess a clinical condition, one difficulty is to evaluate pupillary responses relative to a well-defined baseline. This necessitates defining a period during which no event that could alter pupil size occurs. In research, each study can use an arbitrarily defined baseline, provided it is well characterized in term of luminance and duration, to allow for reproducibility. When considering clinical pupillometry, the conditions defining the baseline must be universal to allow for comparisons between healthy and unhealthy conditions worldwide. The current consensus is to measure pupil size after dark adaptation, for duration lasting from 5 to 20 minutes (Kelbsch et al., 2019). This is very constraining if the goal is to devise tools to be routinely used to assess a clinical condition, as it requires a significant amount of time. In addition, each flash of light inducing a pupillary response must be followed by a period allowing pupil size to return to baseline, in the order of 3 to 10 seconds, although alternative approaches have been proposed (e.g. Maddess et al, 2009, Rukmini et al., 2019).

Unfortunately, dark adapting patients before testing is constraining and long, at odd with the needs in overbooked clinical services. Moreover, obtaining reliable PLR data may requires repeating the test several times, resulting in long lasting sessions.

Dark adaptation may not be necessary with mPFT, because the sustained multipartite stimulation may still reveal the *relative* imbalance of reactivity to light across the visual field, whatever the adapting level. In addition, the mPFT stimulation may be more similar to the ecological situations encountered by individuals in everyday life, where continuous modulation of light intensities is common. To test whether the adapting level deeply changes the mPFT –absolute and relative-spectral power, we performed an experiment (with however a single subject repeating the test many times: N=42 trials, 21 runs per condition) and compared the mPFT power distribution after dark adaptation (10 min., 0-6 lux) or after adapting to day light (4000 to 7000 lux) for 30 to 60 seconds before each run. In the latter case, the mPFT test was ready to launch with a simple click, so that the time elapsed between exposure to day light and the beginning of the test, conducted in a dark room, was less than 30 seconds. We found that the averaged spectral power was indeed slightly different in both conditions, but that the distribution of spectral power across sectors was very similar (Supplementary Figure 7). Some other differences were observed: the lag between the pupillary signal and the stimulus was shorter in the light adapted condition, and the slope of the linear fit computed on pupillary signal was steeper in the light adapted condition, suggesting that adaptation to the stimulus mean luminance developed during the test. The speed of constriction in response to a flash (PLR) recorded at the beginning of a run was also significantly different in the 2 conditions. Although more data should be collected to confirm this preliminary observations, it suggests that dark adaption prior to running the mPFT test may not be mandatory to identify visual defects in patients.

To verify whether our statement that the *relative* spectral power was more stable and reliable across conditions than the *absolute* spectral power (see above and supplementary Figure 5), we compared the *absolute* and *relative* spectral power distributions in the dark-adapted and light-adapted conditions by computing the Pearson’s coefficient correlations between both conditions (dark adapted vs light adapted) for the *absolute* power and for the *relative* power distributions. We found a correlation of r=0.863 (p<0.0001) between the *absolute* power distributions and a correlation r=0.928 (p<0.0001) between the *relative* power distributions, suggesting that, indeed, the *relative* power distribution captures between-sectors regularities not related to the differences of *absolute* power between the 2 conditions.

### Summary of clinical studies done with mPFT

We used mPFT in several clinical studies to evaluate the extent to which it can be used to classify healthy participants from patients (Ajasse et al., 2022; Stelandre et al. 2023; Trinquet et al., in preparation). Acceptability, comfort and feasibility were also evaluated with questionnaires comprising questions about duration, glare, fixating difficulty and fatigue induced by the mPFT test.

In a first study (Ajasse et al., 2022) 3 rare pathologies were tested at the Paris Vision Institute: Retinitis Pigmentosa (N=14), Stargardt disease (N=14) and Leber hereditary optic neuropathy (N=9). Healthy participants (N=14) were also included. In this study, an EyeLink II eye-tracker (SR Research, Ltd, Canada) was used; the stimuli were back projected on a large translucent screen (55.5 x 41.63 degrees of visual angle) while participants sat at 128 cm from the screen. The mPFT stimulus subtended an angle of about 40°.

A second study (Stelandre et al., 2023, article in French) focused on Glaucoma patients (N=28) at different stages of the disease and also involved healthy participants (N=17). In this study, performed in the Ophtalmology Department of Lille, a LiveTrack Lightning eye-tracker (Cambridge Research Systems, Ltd, UK) was used; the stimuli were presented on a computer screen (Dell 210-AXKQ) while participants sat at 80 cm from the screen.

Finally, we used mPFT to test patients with Age-related Macular Degeneration (AMD, N=24), Diabetic Retinopathy (N=29) and Glaucoma (N=8) together with healthy controls (N=40) in a study conducted at the Monticelli Clinic in Marseille (Trinquet et al, in preparation). In this study, a LiveTrack Lightning eye-tracker (Cambridge Research Systems, Ltd, UK) was used; the stimuli were projected on a computer screen (Samsung, SyncMaster 2443) placed in a cabin to control for ambient illumination, while participants sat at 75 cm from the stimulation screen.

For these different studies, performed on different sites, using slightly different setups and protocols (each including other pupillary tests), we computed AUC of ROC, sensitivity and specificity with the fitglm and perfcurve functions available in Matlab (with a binomial distribution and logit link function in the linear regression model), using the power and phase lags of the 9 FOIs. We also used the *relative* power between sectors, computing the ratio of powers of up versus down and left versus right sectors, as well as ratios of peripheral and paracentral sectors and the ratio of the central versus all other sectors (resulting in 12 values). We sometimes also included additional variables (global pupillary variables, see Table 1) derived from the pupillary traces to compute AUC of ROC. Although these other variables do not provide information on the regions susceptible to malfunctioning, they may nevertheless characterize pathological conditions.

When only the spatio-temporal information related to mPFT (FOIs Power and Phase) are included, all AUC of ROC computed for each disease are above 0.9. All AUC of ROC are above 0.95 when more variables are included. In these later cases, the gain in classification is related to the global pupil state variables and to the number of corrections (% Blinks and correction of transients, see Table 1). AUCs of 1 were obtained for both eyes and all diseases in Study 1, 2 and 3, depending on the type and number of variables used for classification. In study 3, pooling the results from all the patients still gives AUC of ROC ∼1, depending on the variables used for classification. Combining the data from the right and left eyes gives somewhat lower values (AUC >0.8).

Although these results are all very encouraging, indicating that pupillary responses elicited with mPFT permit to classify patients and healthy participants with excellent sensitivity and specificity, it must be noted that the number of participants in each study remains relatively small, limiting the conclusions that can be drawn from these studies. Note that more features could be derived from the same data set with no additional cost, between-eyes differences in particular, which could further improve the classification.

Finally, correlations between structural (RNFL obtained with OCT) and functional pupillary responses of mPFT were found in 2 studies (see Ajasse et al., 2022; Stelandre et al., 2023).

Of note: Comparing the mPFT data from the young healthy participants of the present study and the data from study 3 done on a different site (Marseille) with different experimenters, slightly different settings, and with older healthy participants (mean 69, std, see above), give very similar results (Supplementary Figure 8).

### Visual Field, Pupillary Response Fields and structural retinal imaging (OCT)

Do Visual field and Pupillary Response Field probe similar, possibly defective, functional or structural features?

No clear answer to this question is yet available. As RGCs, ipRGCs driving pupillary responses are ganglion cells and may thus be sensitive to harms of the same kind as those targeting RGCs, although their number, functioning and distribution across the retina are different (Zele and Gamlin, 2020). Evidence for a loss of ipRGCs in Glaucoma (Rukmini et al., 2015) reveals pupillary changes proportional to disease severity, but better knowledge of the relationships (in the temporal and spatial domains: degree of temporal and spatial overlap) between the development of visual, structural and pupillary defects would help to characterize them more precisely.

Other studies, in relationship with different and diverse diseases, have reported correlations between pupillary and structural markers in several pathologies (Bremner, Wilhem, Thompson, Rukmini et al., 2019). We ourselves found similar, weak but significant correlations between RNFL and pupillary mPFT power in retinopathies and neuropathies (Ajasse et al., 2022; Stélandre et al., 2023). We did not observe such structural / functional correlation in the population of young healthy participants who performed in the present study. The correlation we do report above when comparing between-eye differences of RNFL and between-eye differences of spectral power deserves more investigations and confirmation as well as a sensible hypothesis to account for it.

Alternately, even if Pupillary Response Fields would carry little resemblance with Visual Fields or structural features seen with OCT, they may nevertheless characterize a functional defect, possibly signaling the existence of a clinical issue specific to the underlying pupillary circuits. Whether or not this malfunctioning entrains, or correlates with, a visual complaint (migraine, glare, over-blinking, blur) needs more work, but several studies found significant correlations between them (Mylius et al., 2003; Harle et al., 2005, Howarth et al., 1993).

### Eye-Trackers and use of mPFT

We have tested mPFT with a few eye-trackers endowed with different characteristics: EyeLink II (SR Research, Canada), Livetrack Lightning (Cambridge Research System, UK), The EyeTribe and SMI Red250 (Sensory Motoric Instrument). mPFT gave good results (clear distinct spectral power peaks at FOIs) with the EyeLink II and the LiveTrack Lightning, but not with the EyeTribe or the Red 250 from SMI. Independently, Mark Wexler in the lab, developed the mPFT stimulus and tested it with a PupilLabs eye-tracker (PupilLabs, https://pupil-labs.com/, Germany), successfully isolating peaks at FOIs in his pupillary response. Informal tests with a Tobii Pro 500 (Tobbi, Ltd, Sweden) and the E(ye)BRAIN eye-tracker develop by Suricog (https://www.suricog.fr/) indicated that pupillary responses also contained significant power at FOIs.

These tests were conducted with commercially available monitors (at 60 Hz) with possibly different sizes and settings and in several lightning conditions or eye-screen distance (hence covering slightly different retinal regions) while still exhibiting well-behaved spectral power at FOIs in healthy individuals.

### Limitations and Future directions

Although the mPFT settings used here elicit reliable pupillary data that are sufficient to perform excellent classification of healthy participants and patients, improvements of the stimulation are possible, and possibly desirable. We already mentioned above the factors that could be adapted (size, luminance modulation amplitude, coupling of TMFs with sectors). One limitation of the present settings is the spatial extent of the stimulus (about 40° of visual angle) that may be too small to detect peripheral visual defects that signal the onset of a disease, as it is often the case for Glaucoma. Simply reducing the Eye/Screen distance would permit to overcome this issue.

Another limitation is the spatial resolution of mPFT, as each sector used herein covers a rather large region of the visual field. We did test a version of mPFT with smaller sectors distributed in 2 opposite quadrants of the visual field, but, although similar classification results were found in Study 1 (Ajasse et al., 2022), two runs are needed to cover the whole visual field, thus lengthening the test duration. However, we noted that visual fields assessed with Standard Automated Perimetry (SAP) may rarely reveal a very focal defect.

A third limitation is the use of an achromatic stimulation, as chromatic pupillometry develops and provides interesting results, related in part to the retinal circuits involving ipRGCs, but is also relevant to characterize visual defects differently altering rods and cones (Zele et al., 2019; Rukmini et al., 2019). We have recently tested a chromatic version of mPFT that provides promising results (Trinquet et al., in preparation).

Finally, the clinical studies summarized in this article involve patients whose pathology is already well characterized, sometimes at an advanced state. Using a pupillary test to follow-up these patients may not be that useful, thanks to the progress of retinal imaging, although it may be relevant to evaluate the effects of treatments on pupillary responses in longitudinal studies. One interesting, but challenging, use of mPFT would be to screen for ophthalmic pathologies in populations at risk, who could develop a still unnoticed pathology, as it is the case in AMD or Glaucoma. To that screening aim, a very large data set and the use of deep-learning appears necessary, and could help determining whether pupillary responses have the potential to early detect and identify retinal or neuronal issues, such as RNFL thinning.

## Conclusion

We presented a novel method to map multifocal Pupillary Response Fields (PRF) in a short amount of time, with little burden for the participants, that permits to distinguish healthy individuals from patients with a variety of neuropathies and retinopathies with excellent sensitivity and specificity.

This method is easy to use, not requiring a dedicated expertise to pass the test, relies on reflexive objective physiological signals, and could be used to complement the current functional examination performed in clinical services (Standard Automated Perimetry, SAP, in particular). Notably, this method can be used without the need of prior dark-adaptation, an interesting feature if it were to be used in clinical settings. In addition mPFT can be used in individuals unable to to understand the instructions to perform a subjective evaluation (SAP), as. elderly with cognitive impairments, children, individuals not mastering the local spoken language, and possibly non-human primates.

Assessing a malfunctioning of the retino-pupillary circuits has interest in itself. It could reveal impairments not seen with SAP, and bring additional information related to the physiopathology of the underlying retinal defects. Moreover, malfunctioning retino-pupillary circuits could account for some of the glare reported by some patients (e.g. glaucomatous patients, for instance, see Hamedani et al., 2021). Indeed, the loss of ipRGCs that occurs in some diseases (Najjar et al., 2021; Gracitelli et al., 2014) degrades the pupillary reactivity to light, which could in turn cause glare in some lighting conditions.

Finally, the use of a sustained stimulation also permits to record and analyze eye-movements instability, as well as blinks, known to also reflect the existence and severity of ophthalmic diseases (e.g. Crabb et al., 2010; Lamirel et al., 2014; Kumar & Chung, 2014).

## Supporting information

Excerpt of Movie of mPFT test

## Acknowledgments

We are very grateful to all the participants who agreed to participate in the different studies presented here. Thanks to Richard Legras who accepted to host the present study, and to the students of the department of Optometry at Orsay University who ran the tests. Thanks to Raphael Barbet who performed data acquisition in some control experiments. We also wish to warmly thank Dr. Frederic Matonti, Laure Trinquet, Frederic Chavane, Dr. Aurélien Stélandre, Prof. Jean-François Rouland, Dr. Catherine Vignal-Clermont, Dr. Cédric Lamirel, and Prof. Antoine Labbé from whom the author learned a wealth of information regarding ophthalmology. Thanks to Benoit Cottereau and Mark Wexler for their relevant advices and suggestions, and to colleagues from the lab for their constructive comments.

## Ethic declaration

All participants gave informed written consent in accordance with the Declaration of Helsinki.

## Data availability

The data are available from the author on reasonable request. The Matlab scripts can be shared with whoever is interested, and the Jeda software used for stimulation is available for download on request.

## Supplementary Figures

**Supplementary Figure 1:**
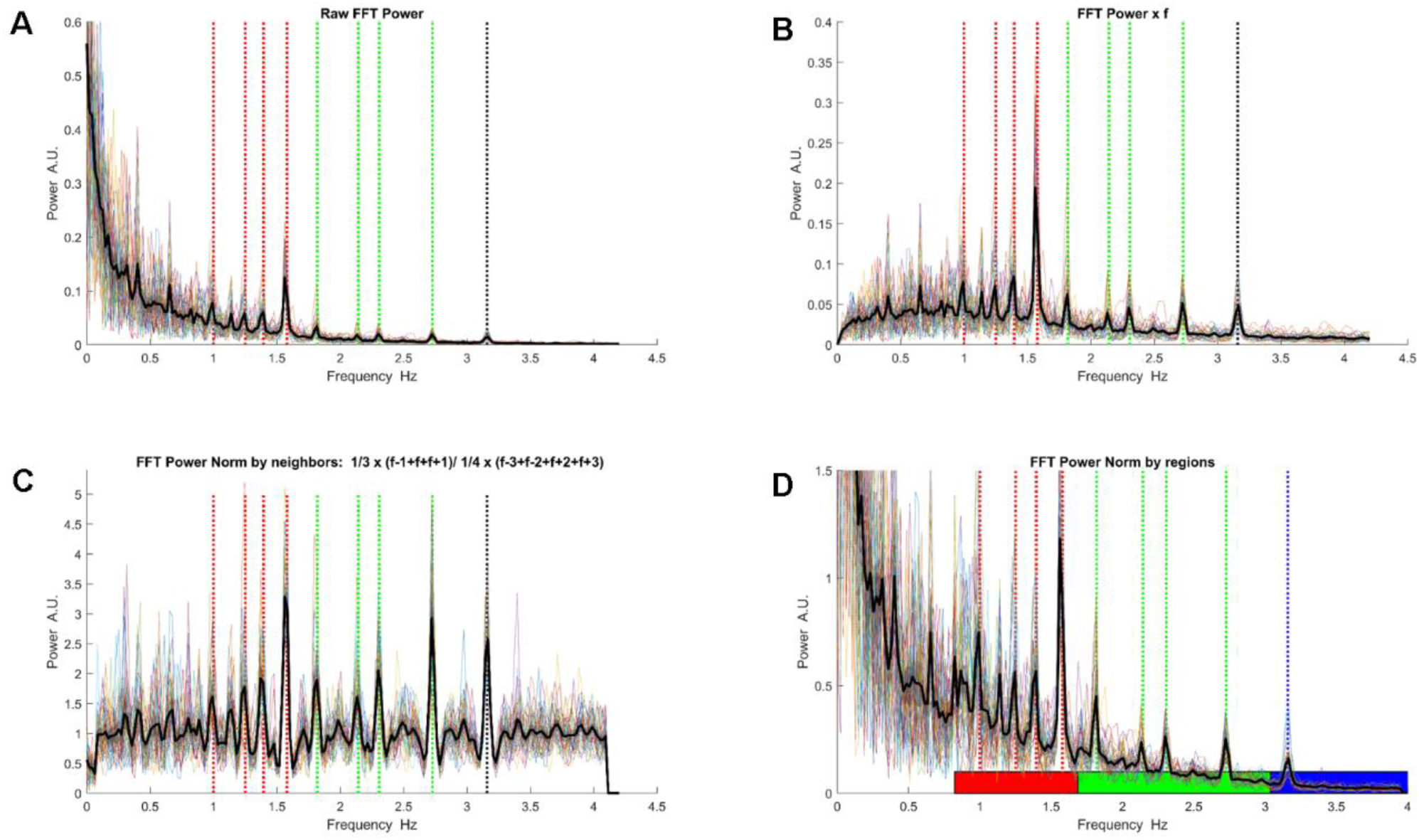
Comparison of different normalization methods. Spectral Power averaged across all participants as a function of Frequency. Vertical dotted lines indicate the FOIs: red lines: peripheral sectors; green lines paracentral sectors; black line: central sector. **A**. Raw FFT spectrum, no normalization. **B.** Normalization by frequency (Power(i) * Frequency(i)). **C.** Normalization by neighbors. Normalized Spectral Power SPn(i)= 1/3x(SP(i- 1)+ SP(i) + SP(i+1)) / 1/4x (SP(i-3)+ SP(i-2) + SP(i+2)+SP(i+3)) **D.** Regional Normalization: The power of each frequency is divided by the mean of all frequencies whose power is less than the mean power in a range. Spectral Power SPn(i)= SP(i)/ mean(SP(j:k)<mean(SP(j:k)) with j and k set for the low frequency range, 0.88:1.74 Hz (red rectangle), the median frequency range, 1.76:3 Hz (green rectangle), and high frequency range (3.02 and 3.94 Hz (black rectangle).

**Supplementary Figure 2:**
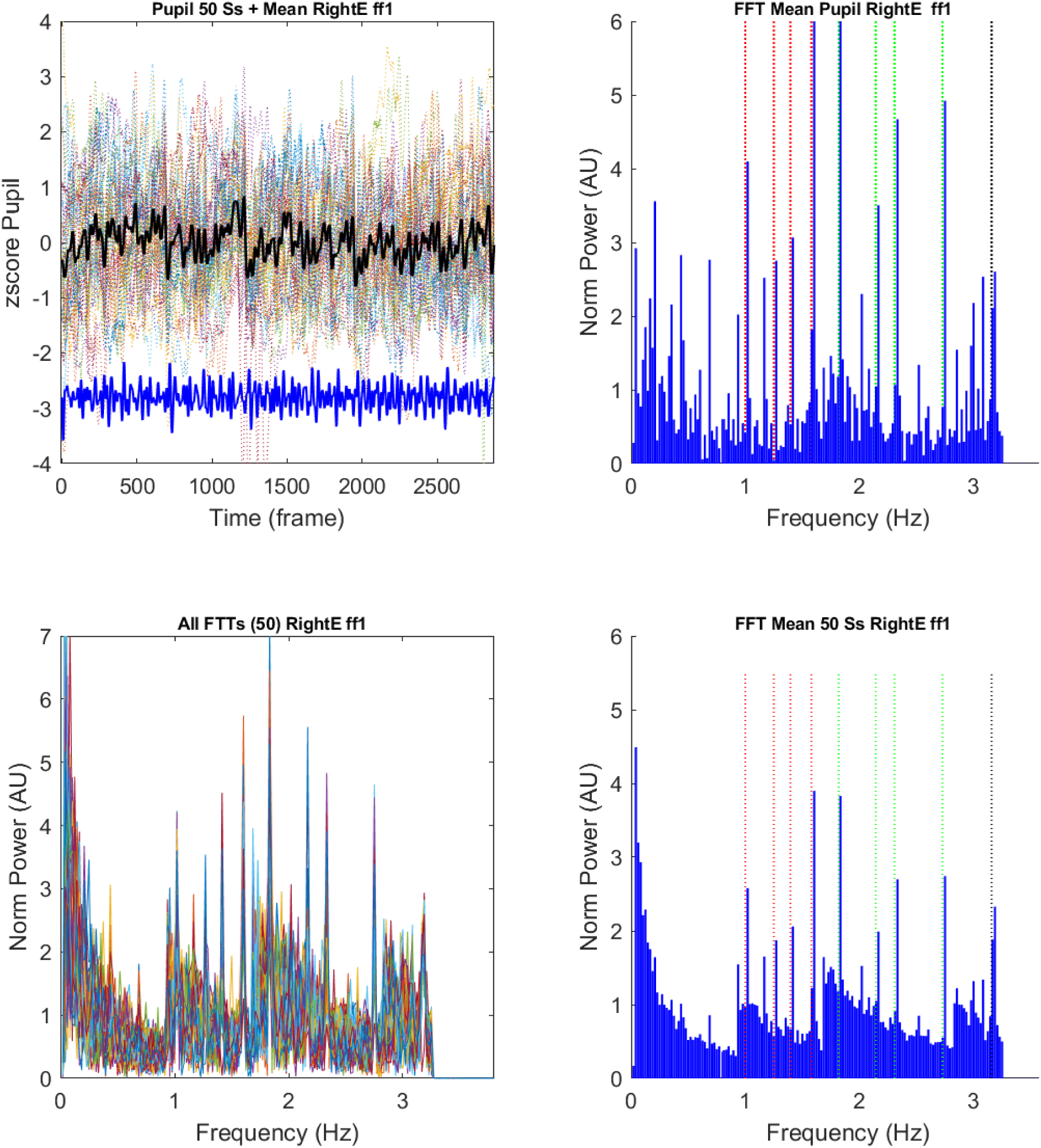
**A.** Top left: pupillary responses during a run for all participants (colored dotted lines), and pupillary response averaged across all participants (black line); Top right: Fast Fourier Transform of the averaged pupillary response **B.** Bottom left: FFTs of all participants; Bottom right: Averaged of the FFT spectra computed for each participant.

**Supplementary Figure 3:**
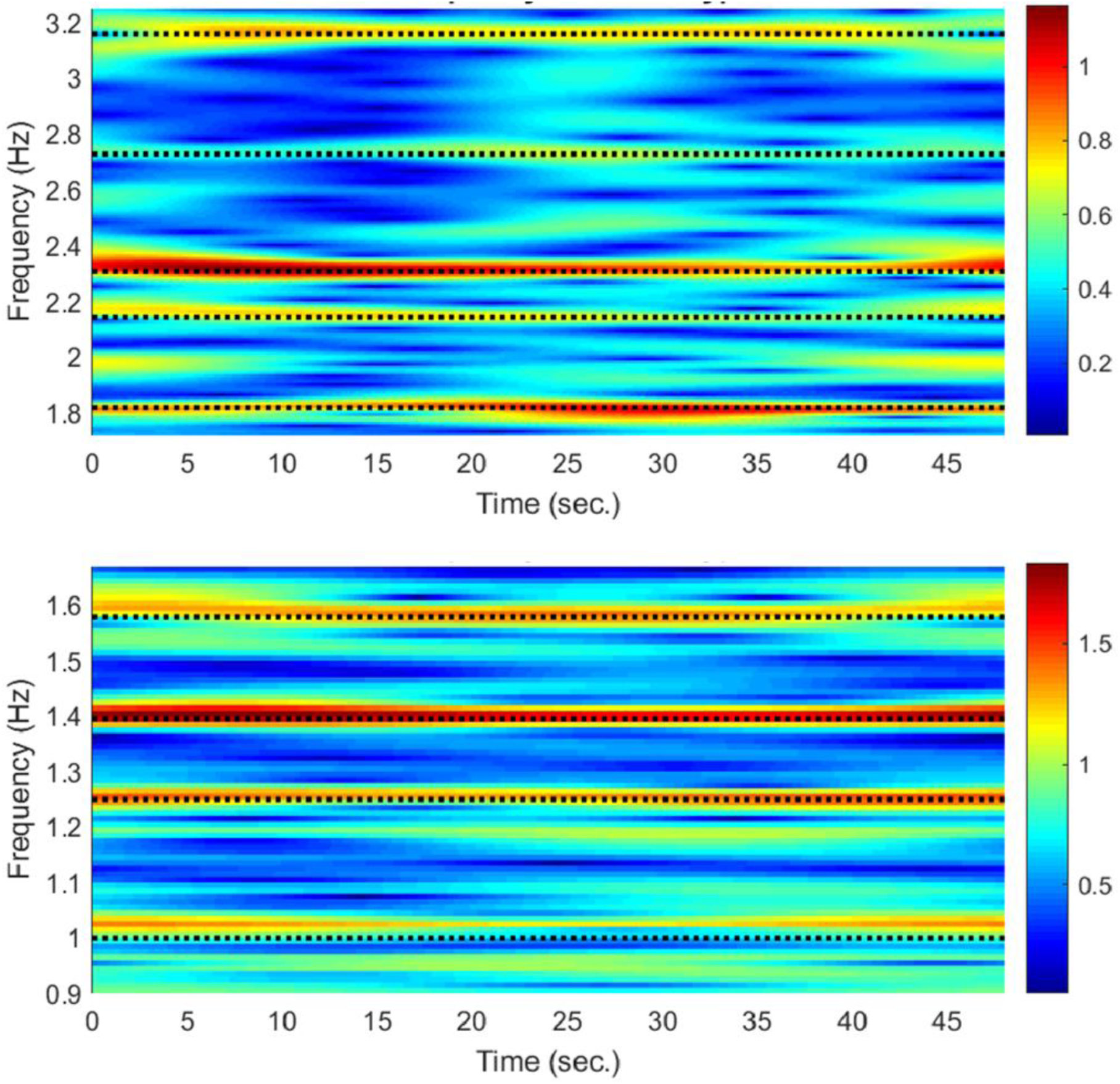
Example of Time Frequency map for a single participant with m=96. Top: Frequencies from 1.81 to 3.17 Hz; Bottom: Frequencies from 1 to 1.58 Hz

**Supplementary Figure 4:**
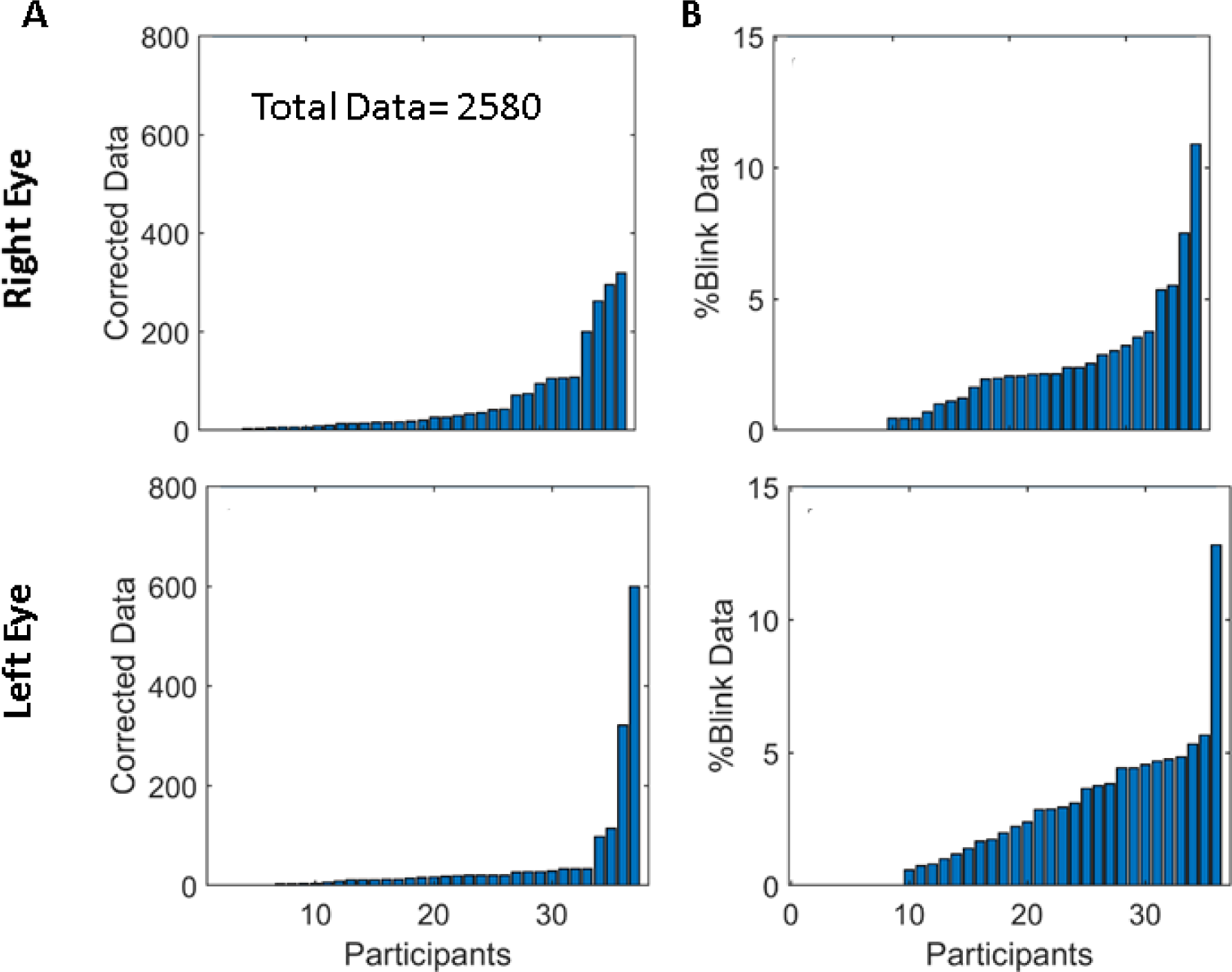
Distributions of the number of corrected data for blinks and transients in the population for the right and left eyes. **A**. Number of data corrected for transients. **B**. Percentage of data corrected for blinks;

**Supplementary Figure 5:**
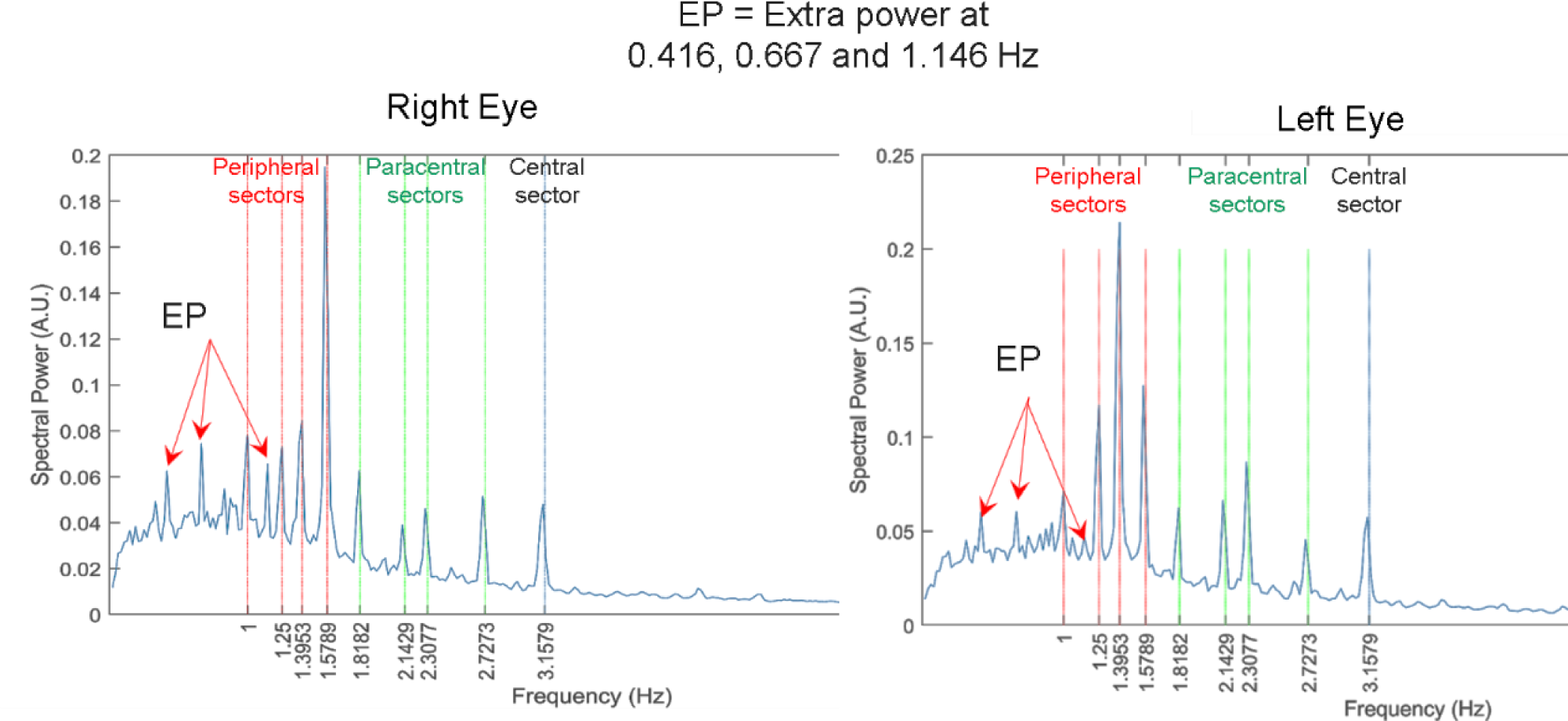
Average of all normalized power spectra from all participants (spectra normalized by frequency: Power * frequency) indicating that power peaks exist at frequencies other than the FOIs, namely at 0.416, 0.667 and 1.146 Hz.

**Supplementary Figure 6:**
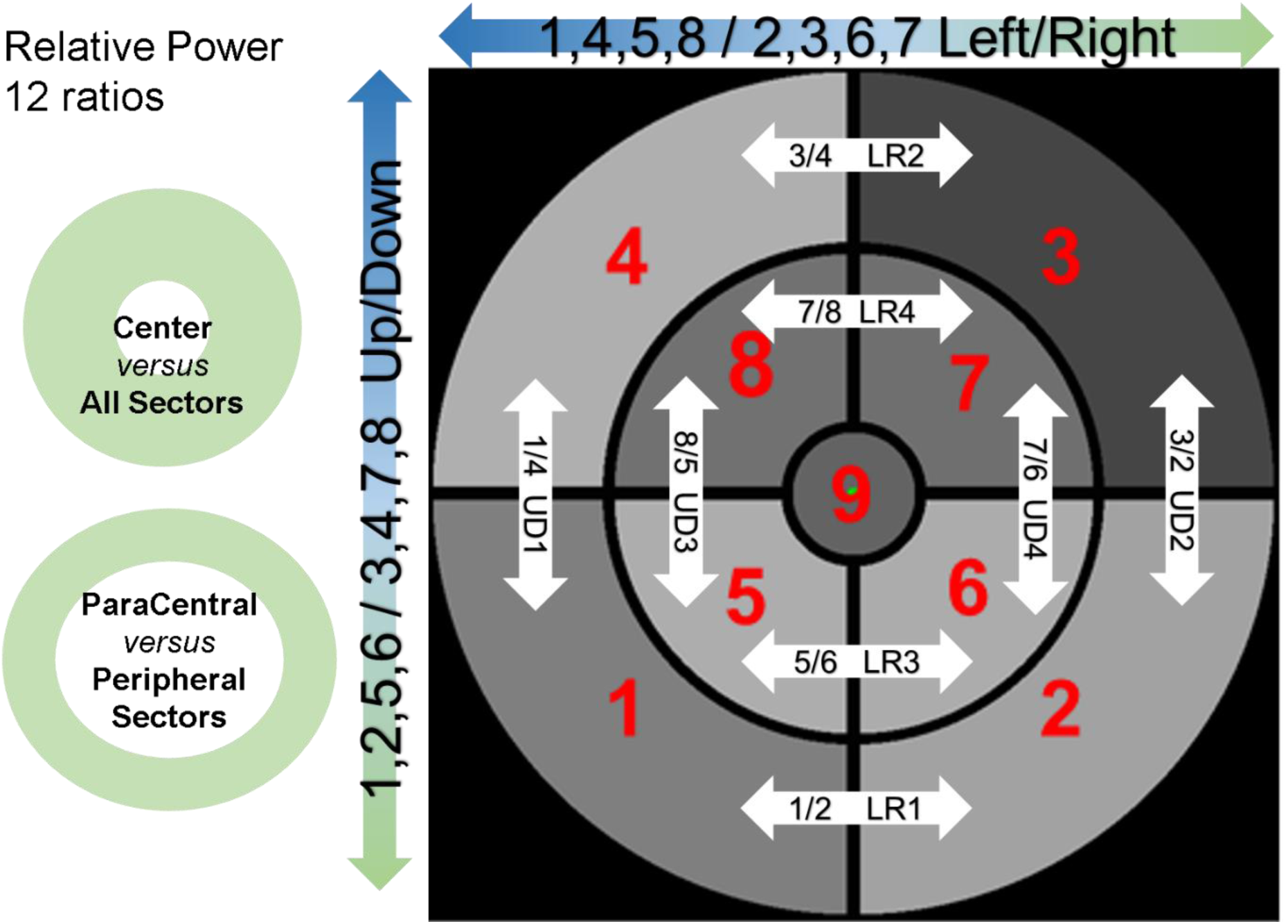
Computation of relative power between sectors: 12 values corresponding to the ratios of power between sectors or groups of sectors are derived from each power distribution of a Pupillary Response Field. These distributions is more robust than absolute power distributions to changes in lightning condition, dark or light adapted conditions, age or medication related to comorbidities that could otherwise alter the overall responsivity of the pupil to light stimulation.

**Supplementary Figure 7:**
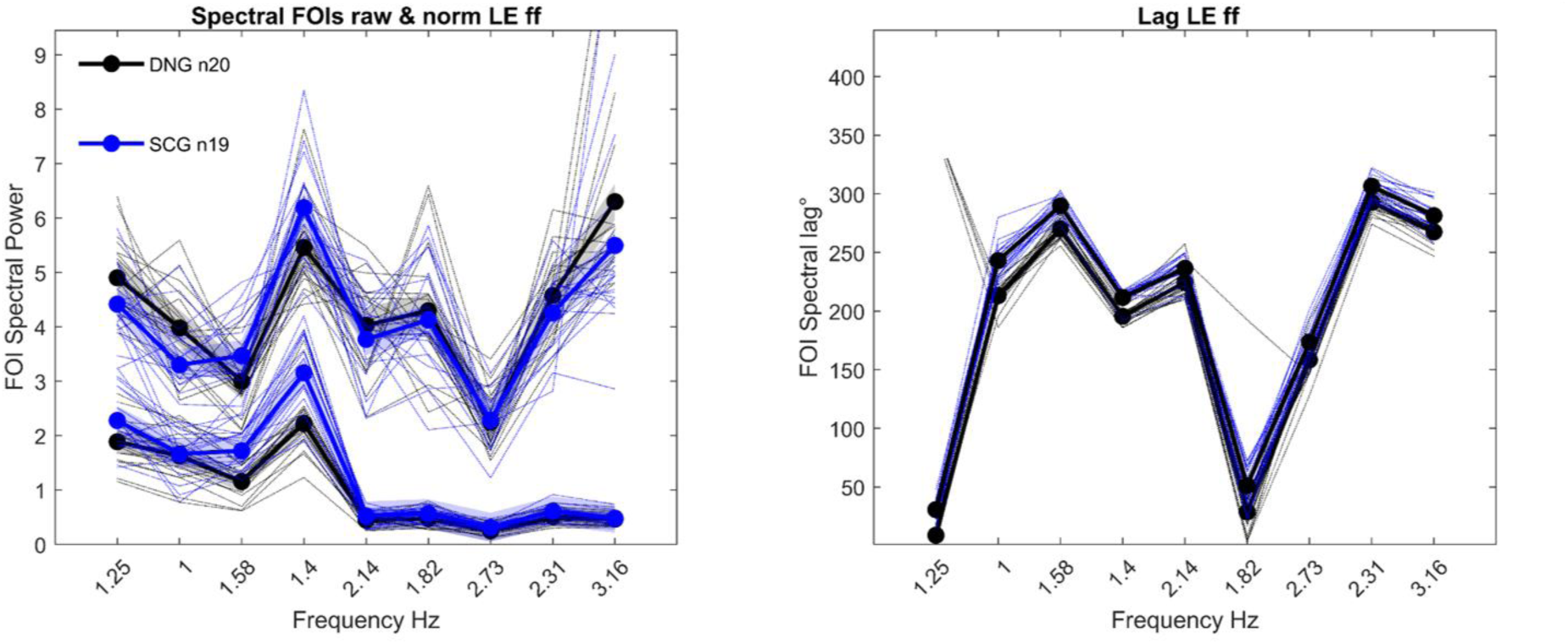
Spectral power at FOIs after 10 minutes of dark adaptation (0-6 lux, black symbols) and 30 seconds of adaptation to day light (4000-7000 lux, blue symbols) before each run in a single subject, but with several trials (N=20 per eye) performed on different days.

**Supplementary Figure 8:**
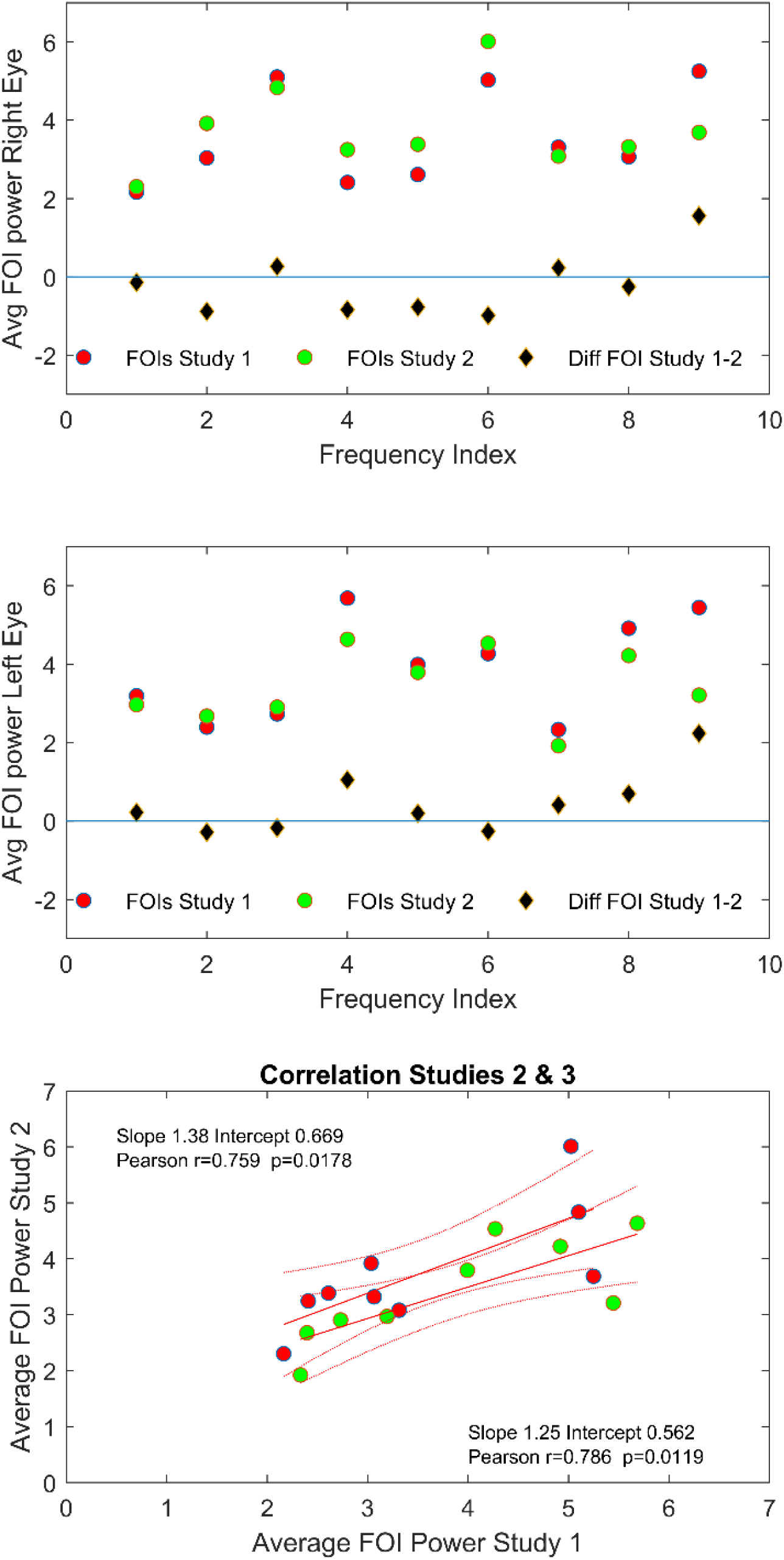
Comparison between the mean FOI powers in the present study (red symbols) and the Marseille Study (green symbols). Top: Distribution of FOI power for the Right Eye in study 1 (red symbols) and study 2 (green symbols) together with the differences of power for each of the 9 FOIs (Black diamonds). Middle, Left Eye; Bottom: Correlation between the averaged FOI Power in the 2 studies. Inserts: Pearson’s correlation coefficients for the right and left eyes.

*Movie #1: mPFT stimulation during a run, starting with a static homogeneous grey stimulus to allow for pupillary adaptation to the mean luminance, followed by luminance modulations. The duration of each phase of the movie has been adapted to limit the movie duration (see Method for a detailed timing of a run)*.

