## Supplementary Figures for "Method to quickly map multifocal Pupillary Response Fields (mPRF) using Frequency Tagging"

**Jean Lorenceau**

**Supplementary Figures**

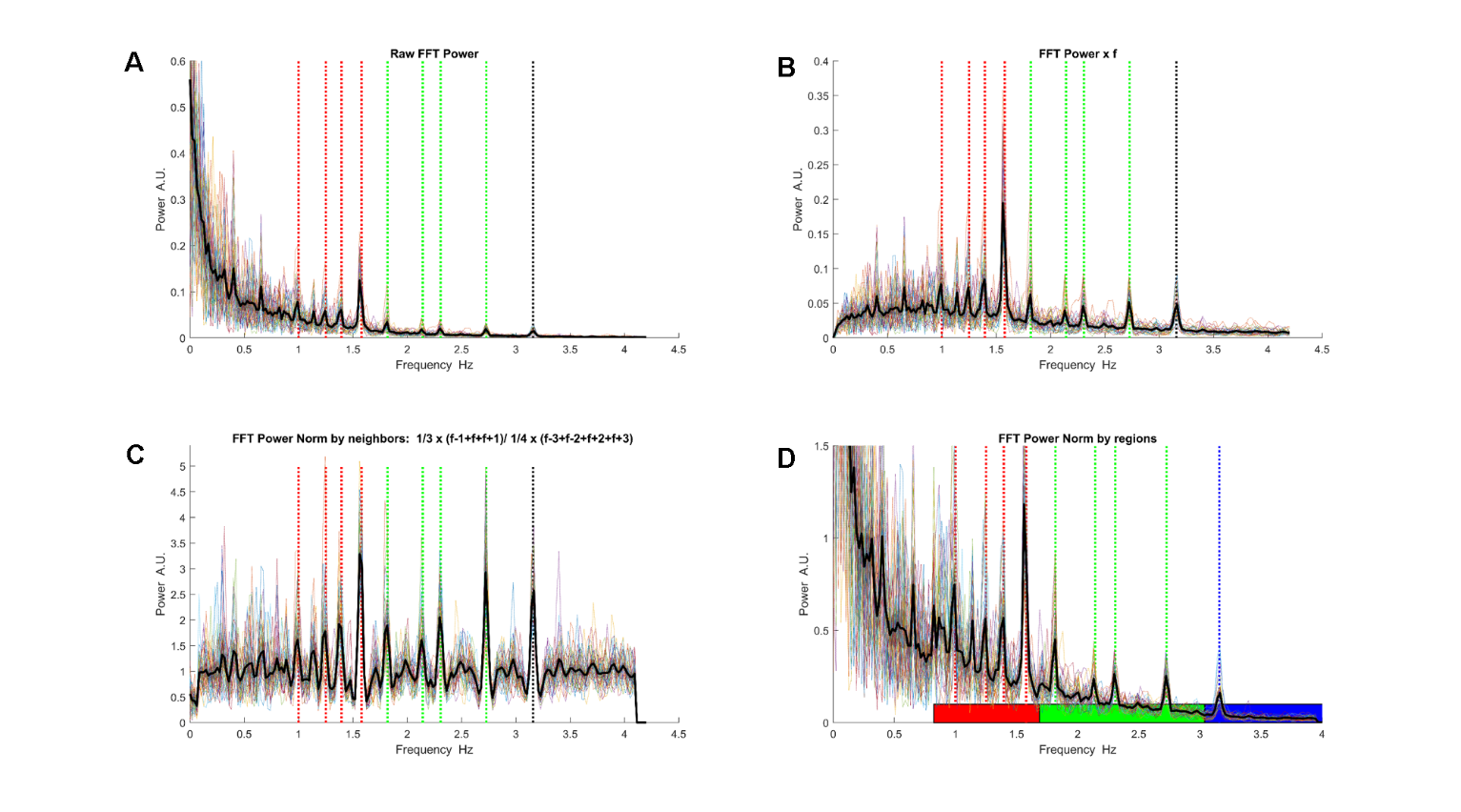

*Supplementary Figure 1: Comparison of different normalization methods. Spectral Power averaged across all participants as a function of Frequency. Vertical dotted lines indicate the FOIs: red lines: peripheral sectors; green lines paracentral sectors; black line: central sector.* ***A****. Raw FFT spectrum, no normalization.* ***B.*** *Normalization by frequency (Power(i) * Frequency(i)) .* ***C.*** *Normalization by neighbors. Normalized Spectral Power SPn(i)= 1/3x(SP(i-1)+ SP(i) + SP(i+1)) / 1/4x( SP(i-3)+ SP(i-2) + SP(i+2)+SP(i+3))* ***D.***  *Regional Normalization: The power of each frequency is divided by the mean of all frequencies whose power is less than the mean power in a range. Spectral Power SPn(i)= SP(i)/ mean(SP(j:k)<mean(SP(j:k)) with j and k set for the low frequency range, 0.88:1.74 Hz (red rectangle), the median frequency range, 1.76:3 Hz (green rectangle), and high frequency range (3.02 and 3.94 Hz (black rectangle).*

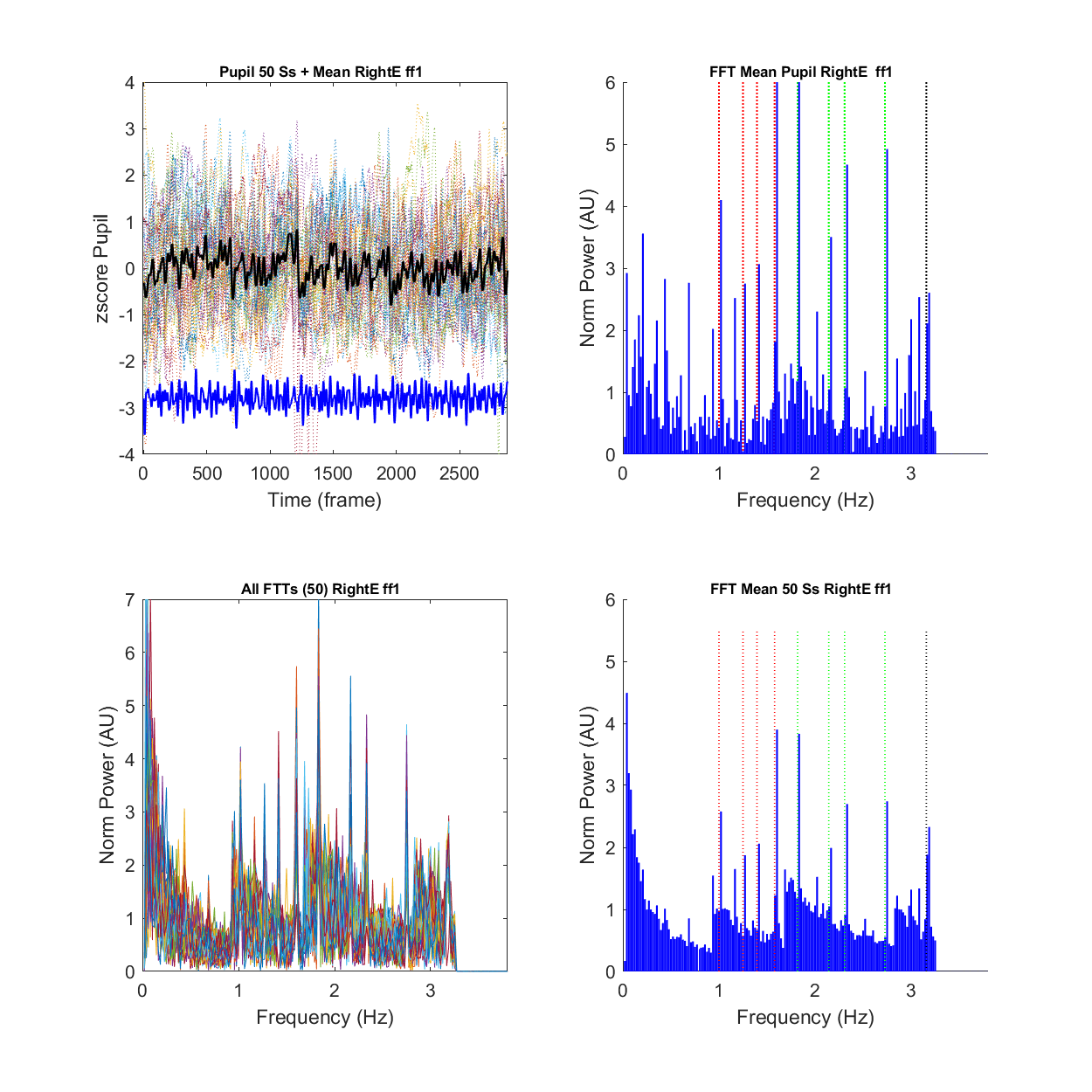

*Supplementary Figure 2:* ***A.*** *Top left: pupillary responses during a run for all participants (colored dotted lines), and pupillary response averaged across all participants (black line); Top right: Fast Fourier Transform of the averaged pupillary response* ***B.***  *Bottom left: FFTs of all participants; Bottom right: Averaged of the FFT spectra computed for each participant.*

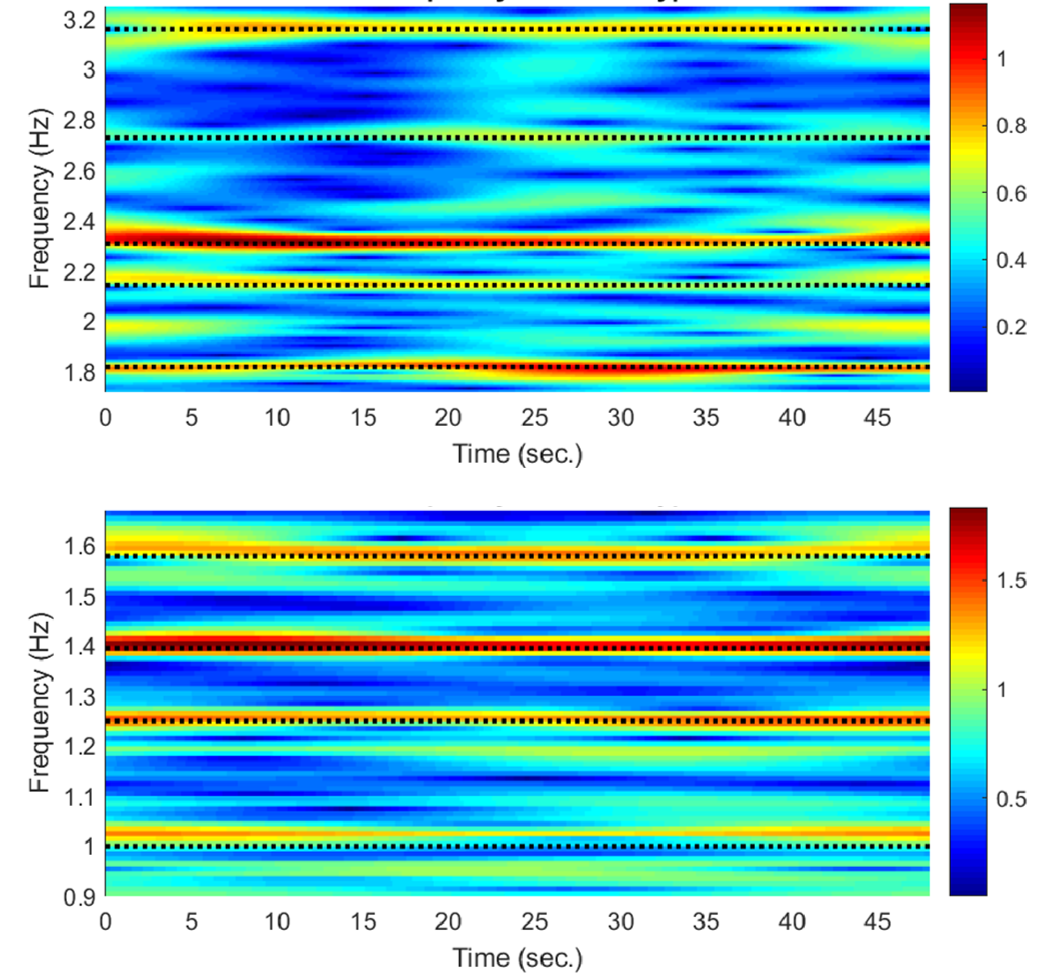

*Supplementary Figure 3: Example of Time Frequency map for a single participant with m=96. Top: Frequencies from 1.81 to 3.17 Hz; Bottom: Frequencies from 1 to 1.58 Hz*

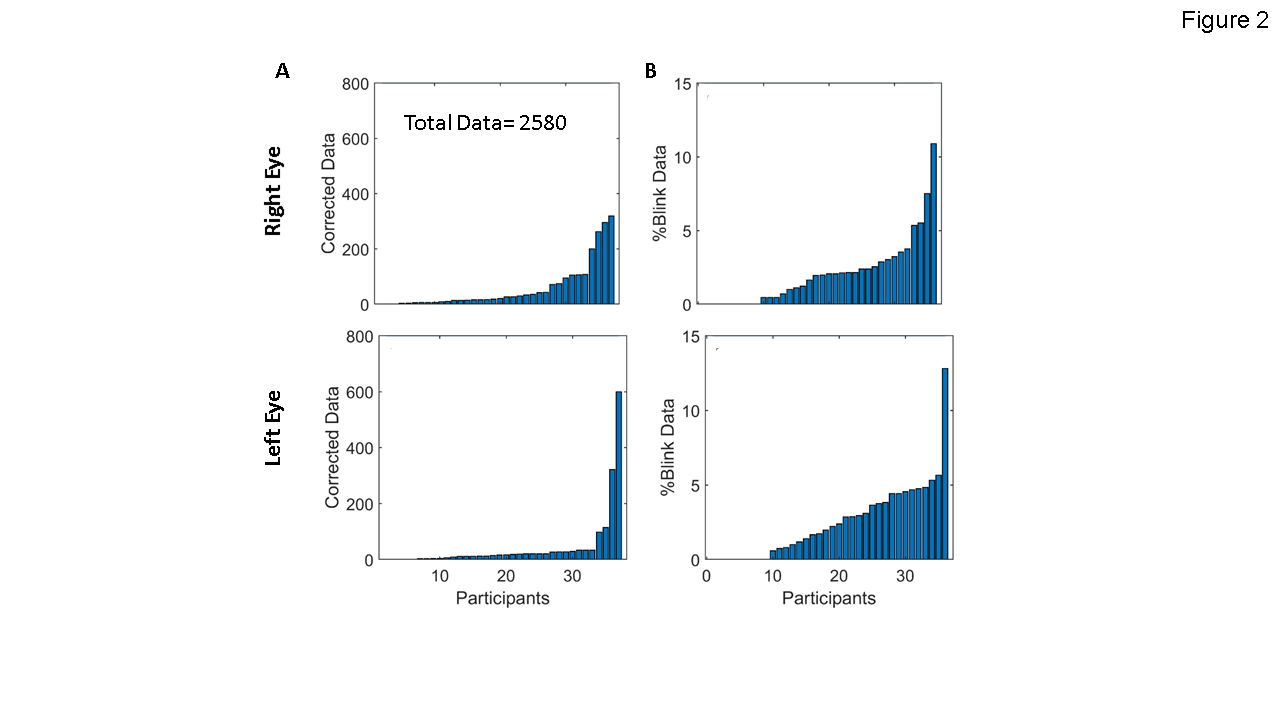

*Supplementary Figure 4: Distributions of the number of corrected data for blinks and transients in the population for the right and left eyes.* ***A****. Number of data corrected for transients.* ***B****. Percentage of data corrected for blinks;*

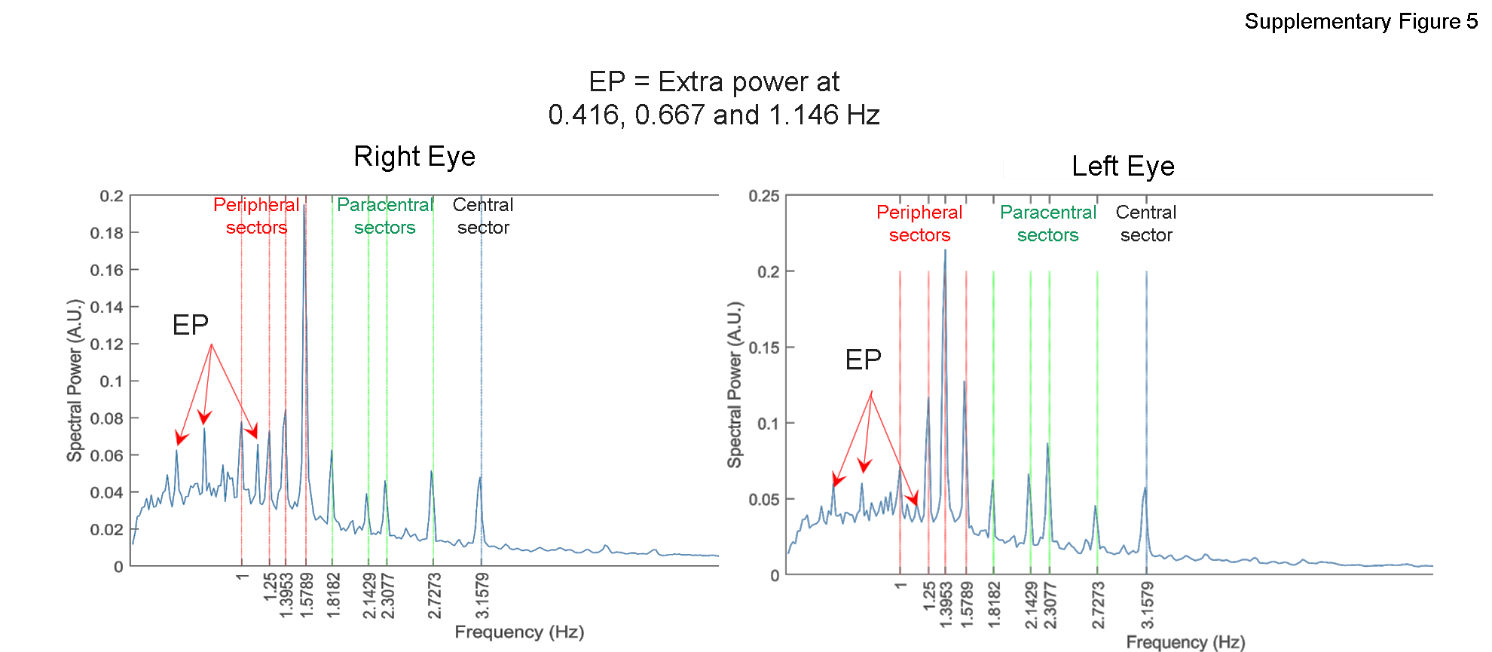

*Supplementary Figure 5: Average of all normalized power spectra from all participants (spectra normalized by frequency: Power * frequency) indicating that power peaks exist at frequencies other than the FOIs, namely at 0.416, 0.667 and 1.146 Hz.*

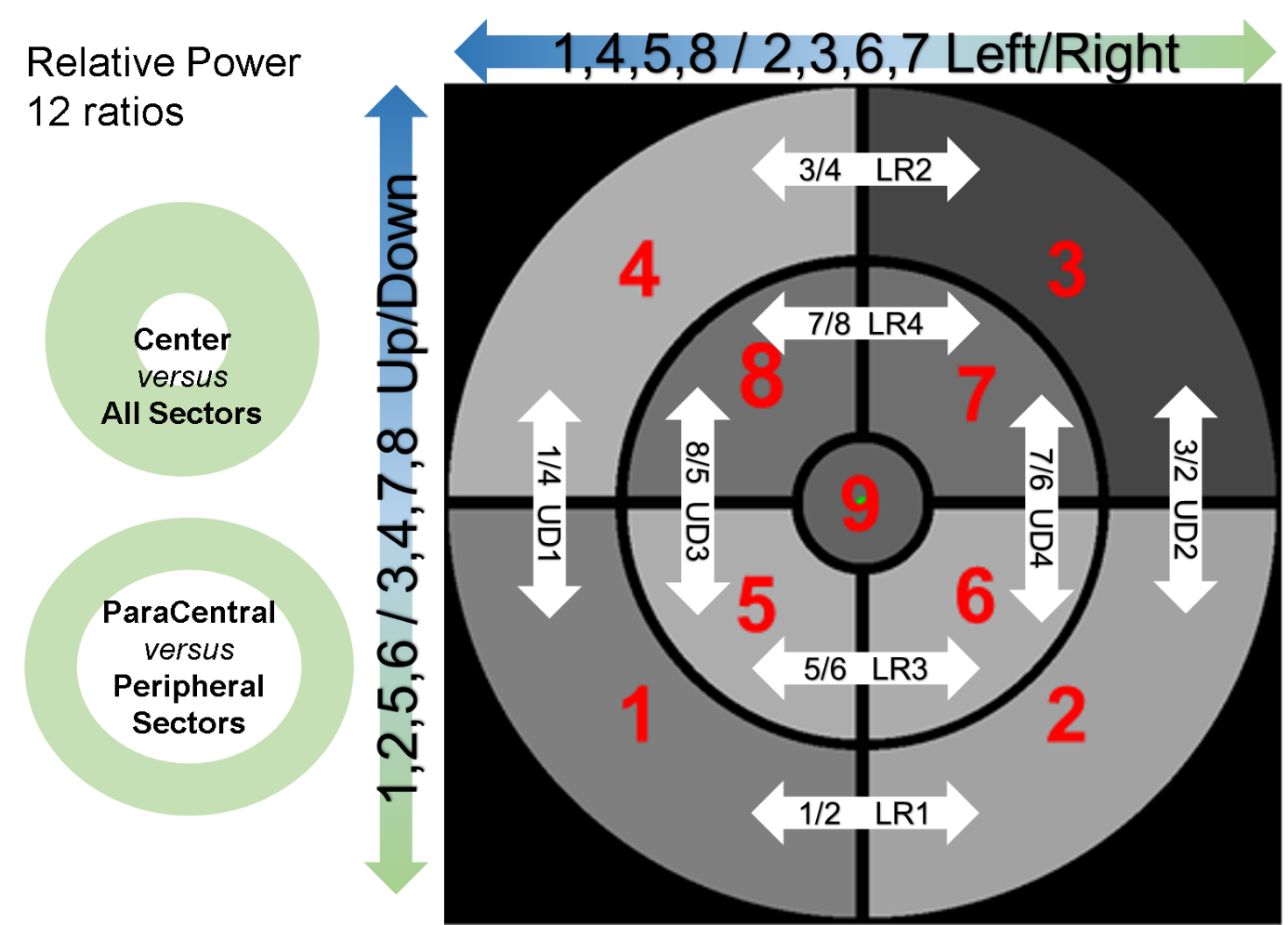

*Supplementary Figure 6: Computation of relative power between sectors: 12 values corresponding to the ratios of power between sectors or groups of sectors are derived from each power distribution of a Pupillary Response Field. These distributions is more robust than absolute power distributions to changes in lightning condition, dark or light adapted conditions, age or medication related to comorbidities that could otherwise alter the overall responsivity of the pupil to light stimulation.*

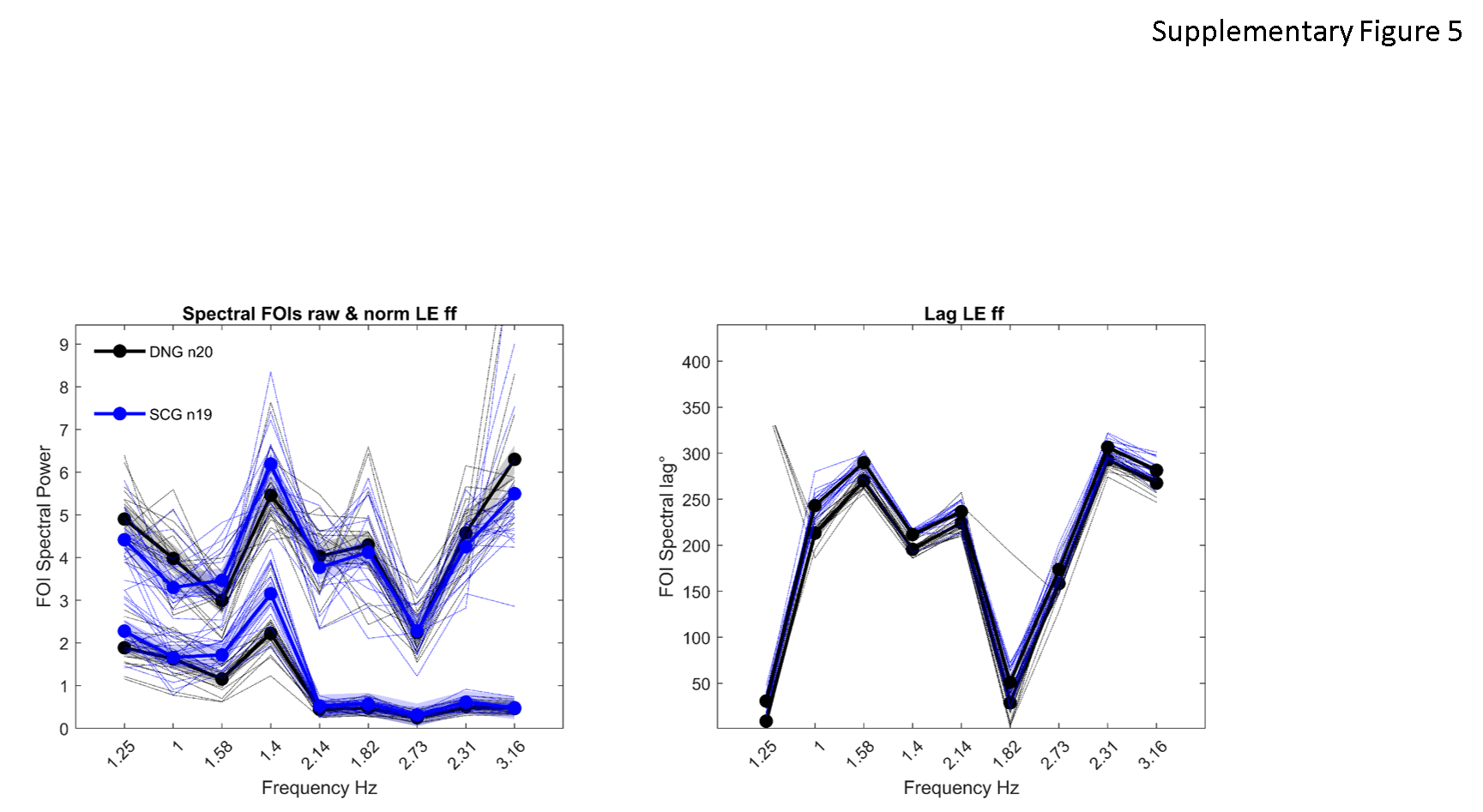

*Supplementary Figure 7: Spectral power at FOIs after 10 minutes of dark adaptation (0-6 lux, black symbols) and 30 seconds of adaptation to day light (4000-7000 lux, blue symbols) before each run in a single subject, but with several trials (N=20 per eye) performed on different days. Raw Spectral Power (bottom lines) and Normalized Spectral power (upper lines) are shown.*

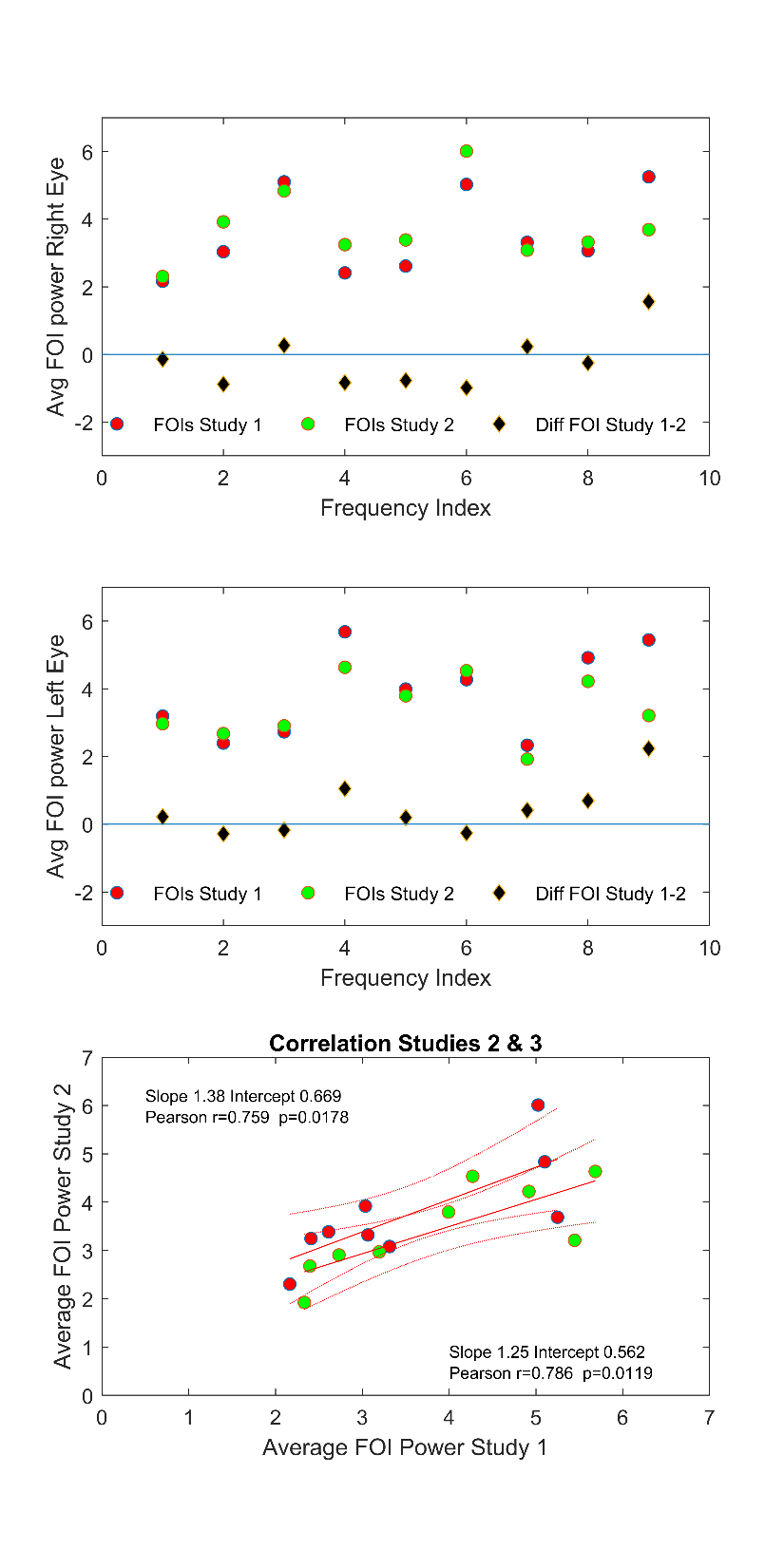

*Supplementary Figure 7: Comparison between the mean FOI powers in the present study (red symbols) and the Marseille Study (green symbols). Top: Distribution of FOI power for the Right Eye in study 1 (red symbols) and study 2 (green symbols) together with the differences of power for each of the 9 FOIs (Black diamonds). Middle, Left Eye; Bottom: Correlation between the averaged FOI Power in the 2 studies. Inserts: Pearson’s correlation coefficients for the right and left eyes.*

*Movie #1: mPFT stimulation during a run, starting with a static homogeneous grey stimulus to allow for pupillary adaptation to the mean luminance, followed by luminance modulations. The duration of each phase of the movie has been adapted to limit the movie duration (see Method for a detailed timing of a run).*
